# Novel topological methods for identifying surprising protein tertiary structure relationships

**DOI:** 10.1101/2023.06.09.544297

**Authors:** Arron Bale, Robert Rambo, Christopher Prior

**Affiliations:** Department of Mathematical Sciences, Durham University, Durham, United Kingdom; Diamond Light Source, Harwell Science and Innovation Campus, Didcot, United Kingdom

## Abstract

We present fast and simple-to-implement measures of the entanglement of protein tertiary structures which are appropriate for highly flexible structure comparison. These quantities are based on the writhing and crossing numbers heavily utilised in DNA topology studies which and which have shown some promising results when applied to proteins recently. Here we show how they can be applied in a novel manner across various scales of the protein’s backbone to identify similar topologies which can be missed by more common RMSD, secondary structure or primary sequence based comparison methods. We derive empirical bounds on the entanglement implied by these measures and show how they can be used to constrain the search space of a protein for solution scattering, a method highly suited to determining the likely structure of proteins in solution where crystal structure or machine learning based predictions often fail to match experimental data. In addition we identify large scale helical geometries present in a large array of proteins, which are consistent across a number of different protein structure types and sequences. This is used in one specific case to demonstrate significant structural similarity between Rossmann fold and TIM Barrel proteins, a link which is potentially significant as attempts to engineer the latter have in the past produced the former. Finally we provide the SWRITHE python notebook to calculate these metrics.

**Author summary:** There is much interest in developing quantitative methods to compare different protein structures or identify common sub-structures across protein families. We present novel methods for studying and comparing protein structures based on the entanglement of their amino-acid backbone and demonstrate a number of their critical properties. First, they are shown to be especially useful in identifying similar protein entanglement for structures which may be seen as distinct via more established methods. Second, by studying the distribution of entanglement across a wide sample of proteins, we show that there exists a minimum expected amount (a lower bound) of entanglement given the protein’s length. This bound is shown to be useful in ensuring realistic predictions from experimental structural determination methods. Third, using fundamental properties of this entanglement measure, we identify two common classes of protein sub-structure. The first are large scale helices, which provide stability to the structure. These helical structures indicate strong structural similarity of two protein families usually regarded as differing significantly. The second class of substructure is one which, though complex, has a small net entanglement. This configuration is physically useful in other disciplines, but its function in proteins is not yet clear. Finally, we provide an interactive python notebook to compute these measures for a given protein.

## Introduction

The sequence of amino acids which form a protein is its *primary* structure and it is always identifiable. Researchers often visualise a proteins global structure via its backbone curve, the discrete 3-dimensional curve whose points represent the central *α*-carbon atom of each amino acid residue, in the form of a ribbon diagram as seen in 1. It has been shown [2] that the specific sequence of amino acids determine the *secondary* structure of a protein. This *secondary* structure represents the shape of local segments of the protein’s backbone curve. The two most clearly defined types of secondary structure are *α*-helices and *β*-strands, represented as helices and flat arrows respectively in 1. When studying a protein’s structure, we are mostly interested in its *tertiary* structure. That is, how the sequence of these secondary structure elements are folded together.

The link between the tertiary structure and functionality of proteins has been well studied. For a thorough review of this area the reader is directed to [3]. To take advantage of this link, structural databases have been compiled, the largest of which is the Protein Data Bank (PDB [4]) which currently contains over 200,000 entries. The advent of machine learning (ML) methods such as AlphaFold [5] and RoseTTAFold [6] has opened the possibility of routinely predicting protein three-dimensional structure. The most recent breakthrough in this area AlphaFold2 was used to predict the structure of over 200 million amino acid sequences deposited in UNIPROT. To better understand these databases, there has been much work in classification of tertiary structures, as well as large subdomain structures, as seen in CATH [7], SCOP [8], and Dali [9]. The families identified in these databases can give greater insight into this correlation between structure and functionality. One such example is the TIM-barrel domain, which is shown to have a consistent binding site across the family [10].

ML methods are trained to identify the relationship between the amino acid sequence and its associated folded structure, using known protein structures available in the PDB as training data. Thus, as any such method can only be as good as the data it is provided as input, low accuracy predictions can be expected for regions not having any homologous sequences in the PDB, or having multiple homologs with widely different structures. Intrinsically disordered regions (IDRs) are a typical example of situations where low-confidence can be expected [13]. IDRs are the most extreme embodiment of a feature typical of any bio molecule: biological function is linked not only to structure, but also to characteristic dynamics. As proteins are flexible molecules, any predicted structure should be considered as a putative representative of a whole conformational space. From a structural comparison standpoint this means there is the need to compare the similarity of structures “up to a wobble”, that is to say characterising whether two structures have a similar overall shape.

A simplified characterisation of this notion is shown in Figures 2(a) and (b). The trefoil curve (a) is knotted in the sense it must be cut in order to turn it into a circle. The curve shown in (b) can obtained by distorting (a) continuously without the curve crossing itself, i.e without any such cutting. Thus in some sense they are folded in a similar fashion, but this would likely not be captured by more standard distance based metrics routinely used for structural comparison. A more pertinent example of real protein structures is shown in 2(c-d) where the smoothed C*^α^* backbones of two proteins given differing CATH classifications, a TIM Barrel (c) and a Rossmann fold (d) can be seen to bear a striking similarity up to a distortion: they have clear helically coiled domains with the same number of coils in each domain and similar relative orientations of these domains. To aid visualisation, the curves shown are sampled every three amino acids so that the local secondary structure geometry does not hide the similarity of these folds.

**Fig 1.**
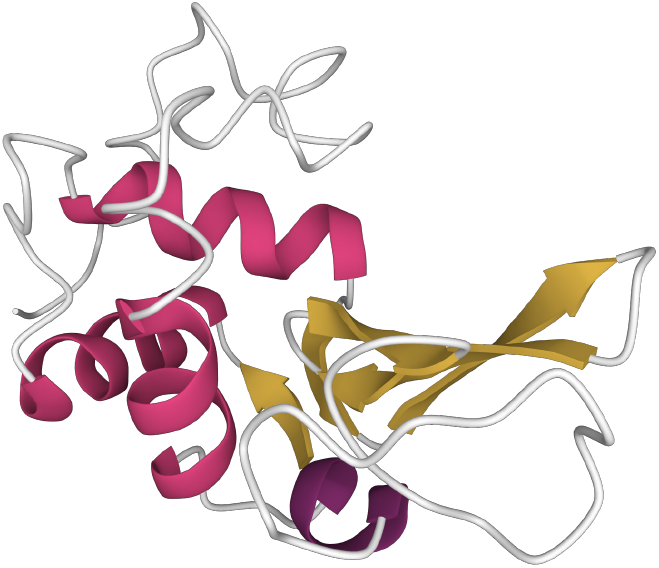
The ribbon diagram for Lysozyme (PDB 1LYZ) [1], with *α*-helices highlighted in pink and *β*-strands in yellow.

**Fig 2.**
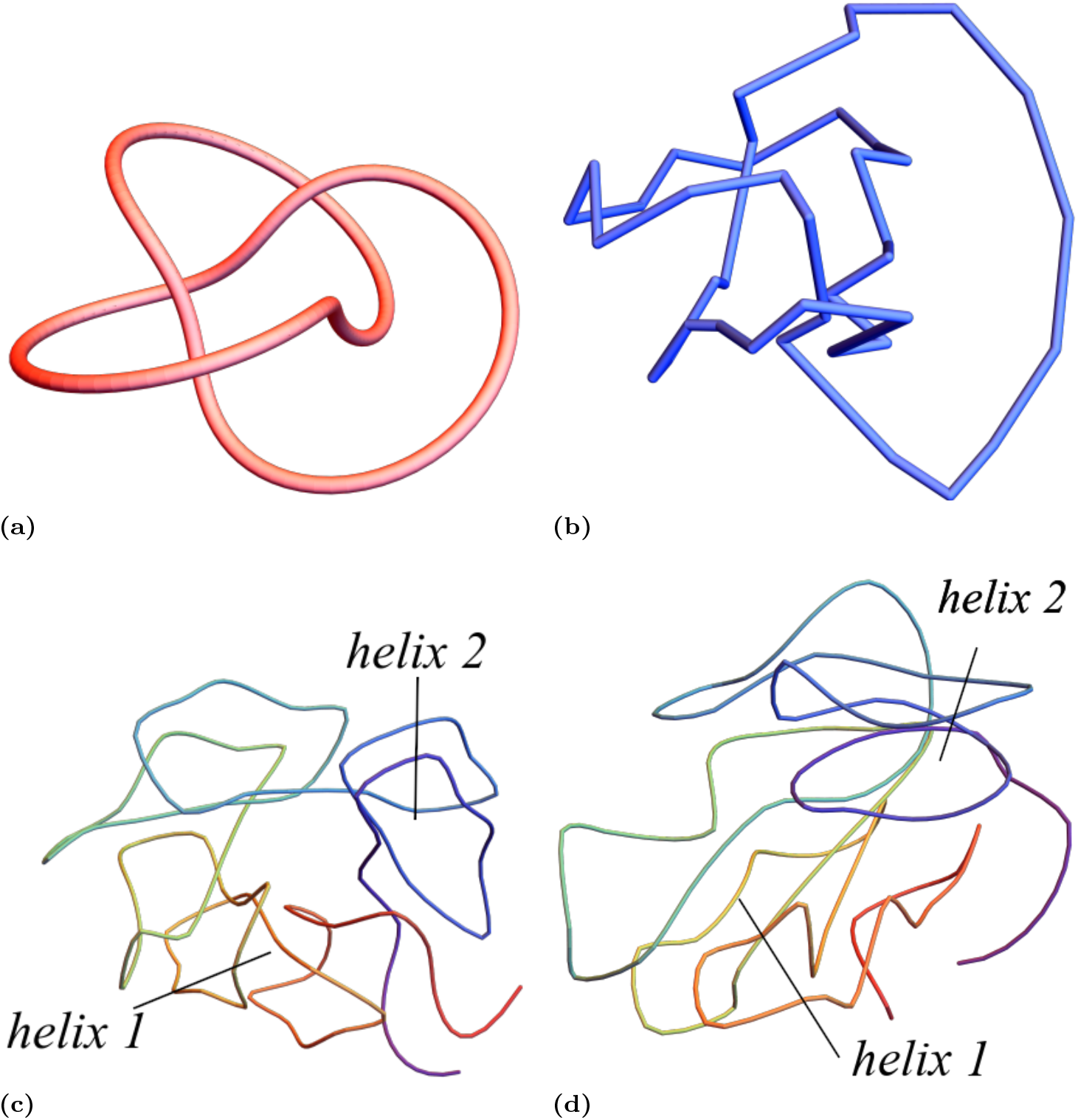
Depictions of the notion of “structural similarity” we seek to quantify in this study. Panel (a) is a trefoil curve: an “idealised fold”, panel (b) is a significant distortion of (a) which preserves its fundamental entanglement. Distance metrics would generally judge the two as significantly different, but topological metrics would judge them significantly similar. Panel (c), a smoothed C*^α^* backbone (with the curve sampled every three amino acids) of the monomer unit of 2-deoxyribose-5-phosphate aldolase (PDB entry 1P1X) [11]. It has two helically wound domains as indicated. The smoothed C*^α^* backbone curve of oxidoreductase, yciK from *E.coli* (PDB entry 3F1L) [12] (with the same smoothing as in (c)) two helical domains are highlighted which are similarly wound as in the structure shown in (c) (but with differing local geometry).

As we will see later, the specific example in 2 is more than just a unique curiosity. Indeed, the results of [14] indicate a very close link between these two domains on the primary sequence, or even evolutionary, level. In [14] the authors used directed evolution techniques to try to design a TIM barrel structure (not the one shown in Figure 2), but found instead a Rossmann-fold-type structure was developed (again, not the one shown in Figure 2). As the following direct quote indicates this was a surprise:

> This surprising result gave us a unique and attractive opportunity to test the state of the art in protein structure prediction, using this artificial protein free of any natural selection. We tested 13 automated webservers for protein structure prediction and found none of them to predict the actual structure.

Both TIM barrels and Rossmann folds have a *β* sandwich structure (anti-parallel strand helices), whose cross-bonds provide stability to the structure. The helical sections of the two figures result from *β*-sandwich motifs. In this study we find this larger scale helical geometry to be highly prevalent in a wide variety of protein structures using the writhe metrics we propose, and further that they are often of a similar scale across a variety of structure types in the CATH classification. We believe searching for similar size helical structures could provide insight into the apparent “mistakes” made in de-novo design methods. This then constitutes a second good reason to develop a new measure of structural similarity, in addition to the need to handle structural flexibility.

A third motivation for developing a more flexible notion of structural similarity is that is is vital for one of the most common techniques used to study the tertiary structure of proteins: Biological Small Angle X-ray Scattering [15–19] (BioSAXS). BioSAXS can be used for structures which fail to crystallise or that are too large for NMR, *e.g* in [20] the structure of the uncrystallisable retinoic acid receptors (RAR-RXR) is determined via SAXS experiments. It is also an ideal technique to study the structure of proteins in solution and often indicates the obtained crystal structure or computationally predicted structure is not the same as that adopted in solution [21, 22]. The cost of this technique however is the difficulty in interpreting BioSAXS data, with the random motion of molecules in solution leading to isotropy of scattering pattern. This limits the available information directly from experiment to only the distribution of pair-wise molecular distances (no orientational information) [16]. Additionally the flexible dynamics of many structures in solution means the method’s spatial resolution is typically limited to about 7.5Å (as quantified by the Shannon sampling theorem [23]). As indicated in [24] RMSD comparisons of ab-initio predictions at this level of accuracy cannot distinguish significantly different geometries of the protein backbone where a topological based method could (similar to but less sophisticated than the one proposed here). We shall demonstrate that the writhe based metric introduced in this study can be used to limit predictions from BioSAXS data which are unrealistically folded.

So our aim in this study is to develop structural comparison metrics which can answer questions such as can one discrete curve be turned into another without significantly folding/unfolding its tertiary geometry? These metrics should, for example, class the pairs (a)-(b) and (c)-(d) from Figure 2 as markedly similar. The most commonly used methods of structural comparison are based on root mean squared comparisons [25]. Two structures are rotated such that the sum of the minimum Euclidean distance between sequentially aligned points is minimised, the so-called root-mean-squared distance (RMSD), very often the comparison is between similar subsections (for example one can compare hinging proteins this way). There are various ways of making this comparison and it is complicated by the fact the alignment of the proteins itself is not necessarily a uniquely well defined problem. Another measure of protein similarity is the TM-score [26], given by

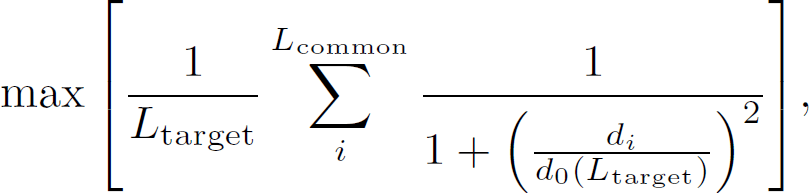

where *L*_target_ is the length of the target protein, *L*_common_ is the number of residues in common between the template and target proteins, *d_i_* the distance between the *i*^th^ pair of residues, and *d*_0_(*L*_target_) a distance normalisation term. By having the *d_i_* in the denominator of the sum, the TM-score is less sensitive to local structural errors than the RMSD, and is therefore seen as a better measure of global similarity. The TM-score falls between 0 and 1, with scores higher than 0.5 being seen as having roughly the same fold.

We will quickly run through a few of the readily available alignment methods for the two proteins in Figure 2(c)-(d) to show how they miss the similarity that is visually apparent. The iPBA method proposed in [27] is an alignment method based on Protein Blocks (PBs), an alphabet of local protein structures. This method gives a few colour coded outputs for the quality of alignment, all of which are bright red in this case, so no indication that these two proteins are similar. Alignment via the DALI method [9] gives a poor RMSD of 4.7Å. The TM-align server [28] gives an RMSD of 5.89Å and TM-score [26] of 0.39, again indicating no real structural similarity. The FATCAT method [29] uses a more flexible approach to alignment, allowing a certain number of twists in the backbone to align residues. This method gives a better RMSD of 3.1Å, but a TM-score of 0.24, again indicative of a lack of structural similarity. This is not a bad thing, the fact the two proteins are classed as different in the CATH results form differing cross-connectivities of their strands helices, which is why the helical geometries are locally somewhat different (the Rossmann-helices are visually wider); nonetheless there are the same number of helical turns in the structure. All of this is to say that the usual alignment based distance metrics, despite their many merits, are not suited to the types of similarity we are aiming to quantify in this study.

These structure comparison methods are also not best equipped to compare tertiary structure models derived from BioSAXS data. For most SAXS data, the resolution will not be able to pinpoint the exact location of each residue. So, we need structural measures which are able to give information about the folding type of the protein without suffering from the noise of small scale changes. As we pass predictions into molecular dynamics simulations, provided the overall fold is correct, the small scale errors will relax into the correct structure so we need to try to identify predictions which are likely to relax to the same state. So we turn to an alternative metric class: topological measures.

### Topological metrics

It is worth highlighting a simple example of the pathology one can encounter when utilising such distance/deformation metrics to motivate the use of topological metrics. Consider for example the two figure 8 curves shown in 3. These curves differ only by the relative orientation of the crossing. The RMSD between these two conformations, as measured for the discretisation seen in 3(c), would be very small (as demonstrated in the supplement of [24]). This is because one could move from one conformation to the other by moving only a very small subset of points. This change however is not realistic, it would require cutting the curve and passing it through itself (the breaking of bonds). The writhe based metrics we introduce here classify these two structures as significantly different and they are from a family of metrics called topological metrics.

Topological methods are those derived from aspects of Knot theory [30, 31], of which our method can be said to belong. Topological metrics are invariant to rotations and translations and **do not require alignment**. A second potential advantage is that they classify structures up to isotopy, *i.e.* they measure two structures as similar if they can be distorted into each other **without having to construct the distortion**. This would mean for example a disordered protein’s structure would likely always be classed as the same/very similar. As shown in [24] they classify the two figure 8 curves in Figure 3 as significantly different and as we shall see in section classify the pairs (a),(b) and (c),(d) from Figure 2 as significantly similar.

**Fig 3.**
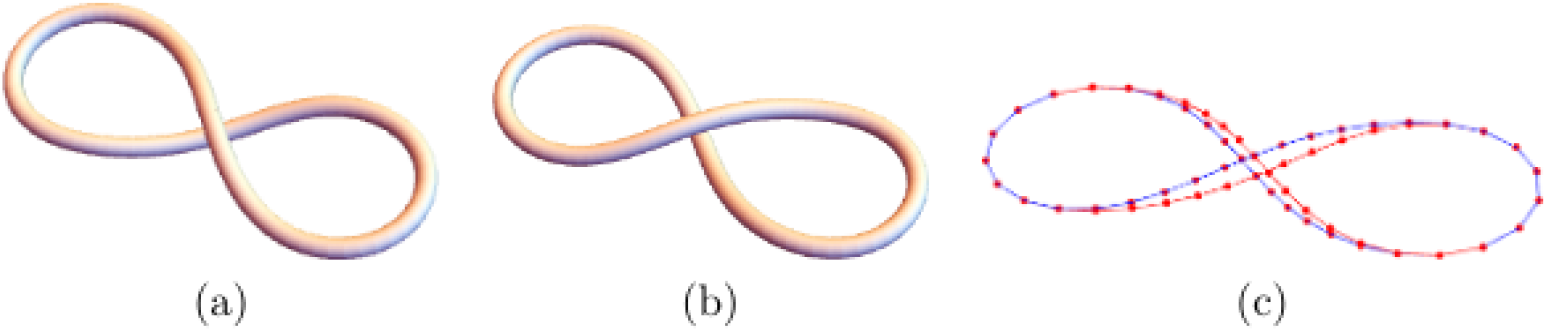
In (a) and (b) we see two figure 8 curves, which differ only by the orientation of the crossing. In (c) we see a discretisation of these two curves, from which an RMSD score can be calculated.

One difficulty with implementing topological measures of protein structures is the fact that proteins are open-ended curves. Mathematically a knot is an embedding of the circle in 3D space which allows for classification of curves which can only be reached by crossing themselves. To overcome this one can “close” a protein by projecting two parallel lines from the protein’s endpoints to a point on a sphere surrounding the backbone curve. To this closed curve we can then assign a knot type. Since these closure lines pass by other sections of the protein, introducing entanglement, we perform this closure for many projections to find the most commonly occurring knot type. Knotting in proteins has been shown to be conserved through evolution [32], however, the question of how and why proteins form knots remains open. On the one hand, knots create a barrier to efficient unfolding as seen in [33]. On the other hand, knots can provide extra stability to proteins [34], a functional advantage that could hint at their evolutionary persistence. To add to their intrigue, knotted structures are in fact extremely rare in proteins [35]. To study knotting on various scales and subdomains, a knot fingerprint was introduced in [36]. The knot fingerprint is a lower triangular matrix, where the (*ij*)^th^ entry is the knot type of the subsection of the protein between the *i*^th^ and *j*^th^ amino acid.

A major issue with this method from our perspective is the aforementioned rarity of knotting in proteins, making it hard to differentiate structures, even when considering the substructures via the fingerprint. In [24] an alternative that considered the distribution of the second most common knots was shown to differentiate proteins which were not knotted, but this metric was far too unwieldy and hard to interpret. There have been some other extensions of the knot theory approach, for example with the introduction of knotoids. Knotoids are essentially a generalisation of mathematical knots, allowing for distinct endpoints [37]. With this generalisation, the world of knotoids is richer than that of knots, and therefore better able to differentiate protein structures that classical knot theory approaches may miss, as in [38]. A knotoid based approach to protein folding problems has also been explored, in [39] the authors are able to uncover a novel folding pathway for shallow knotted Carbonic Anhydrases. Though this approach has been shown to address some of the shortcomings of the classical knot based approach, it is still not best suited to our purposes. Most significantly, knot(oid) classification algorithms are computationally expensive. To speed up these calculations, the backbone is reduced to a minimal representation via the KMT algorithm [40], and then the knot type of this simplified curve is determined. However, this approach loses some information about the specific arrangement of secondary structures, which is vital for our purposes. For example, knotoids are not designed to detect the similar helical geometries we see in Figure 2(c-d). So we turn to writhe measures which include some of these topological properties but also have their own unique properties which suit our purpose

### The writhe and its use characterising proteins

The formal mathematical definition of the writhe of a three-dimensional curve **x** with tangent vectors **T** is given by the Gauss linking integral

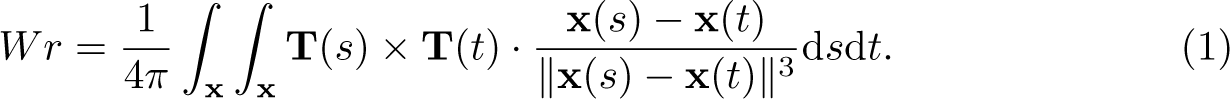

As proteins are discrete curves whose points represent the C-alpha atoms, we use the discrete analogue of the writhe as given in [41]. An intuitive understanding of the writhe calculation is as follows. In Figure 4(a) we see a trefoil curve viewed from a given direction. One can count four crossings of this curve from this perspective and they are given a sign as indicated in Figures 4(c) and (d). The signed sum of these crossings is –4. It is an (oriented) measure of the amount the curve wraps around itself. In panel (b) we see the **same** curve viewed from another angle, there are now only three crossings visible, again with the negative sign. It can be shown [42] that the formula 1 is the average of this crossing number taken over all possible view points, so it measures mean entanglement. The writhe of the figure 8 curves shown in 3 are respectively ≈+1 and –1, so the structures are classed as fundamentally different.

**Fig 4.**
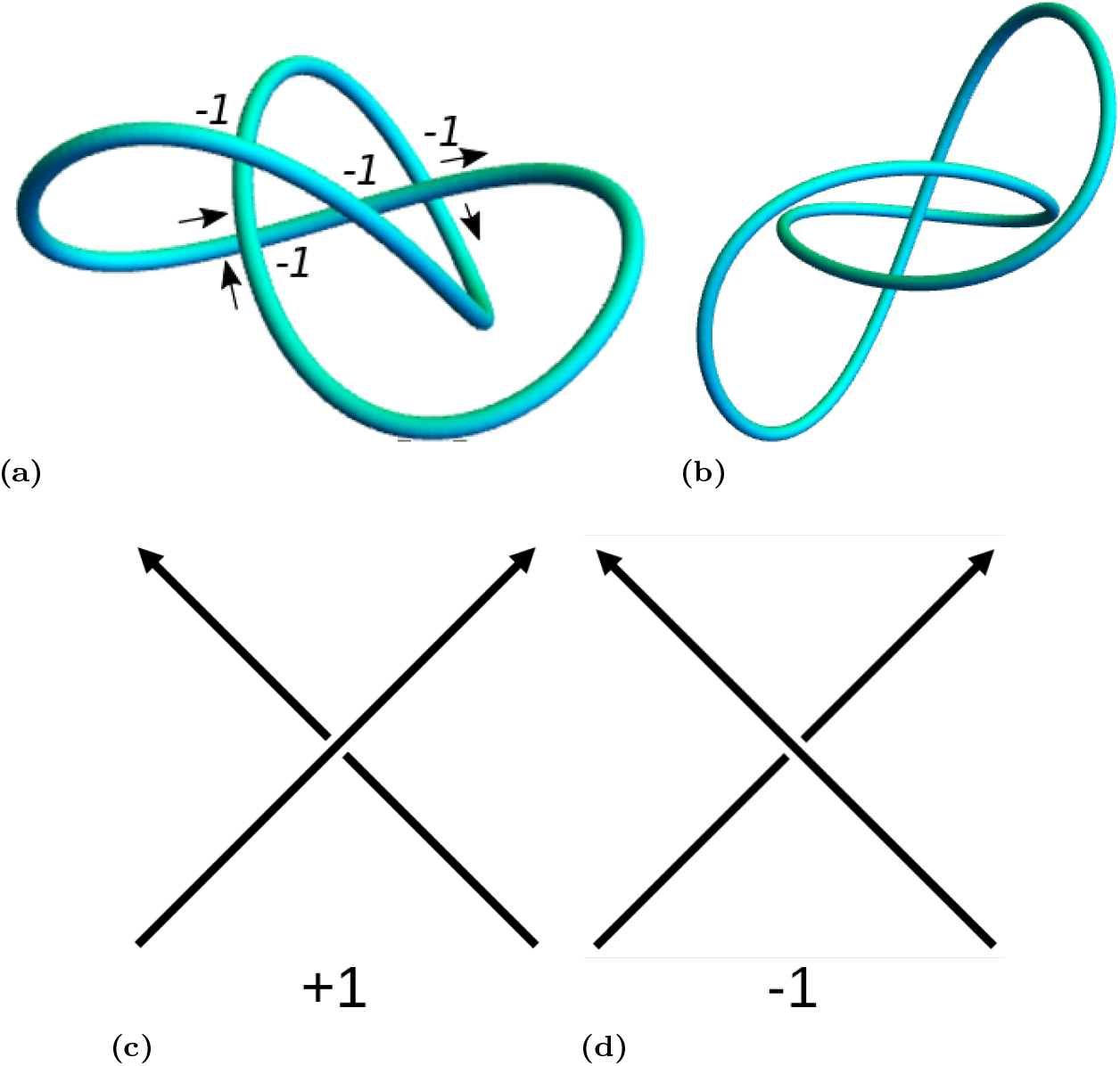
Intuitive interpretation of the writhe. Panel (a), we see a trefoil curve viewed from a particular angle and note the crossings. They are given a sign based on the rules show in (c) and (d), and the total sum (-4) is taken here. In (b) we see the same curve form a different angle and only see three crossings here (-3). The writhe is an average over all projections.

The writhe has long been used to quantify the topological behaviour of DNA. Studies show strong links between the biological function of DNA and its topology, both as a critical role in constraining the elastic energy of [43–48] but also to quantify the role of DNA complexity in various biological processes. For example, in [49] the topological complexity of the DNA as measured by its knot type plays a key role in the rate of DNA ejection for Bacteriophages. In [50] it was demonstrated, using cryo-electron microscopy, that a decrease of DNA-DNA repulsion by increasing concentrations of counter ions causes a higher fraction of the linking number deficit to be partitioned into writhe. In [51] it was shown that DNA molecules from bacteriophage P4 were highly knotted. The use of both experimental and computational modelling techniques indicated a tendency towards high-writhe configurations. In particular it was demonstrated that DNA knots cannot be obtained by confinement alone but must include writhe bias in the conformation sampling. This is just a small subset of the research on this topic. For the interested reader a far more comprehensive review of DNA topology from a mathematical perspective was conducted by [52]. Finally [53] used a combination of both local and non local writhe based metrics to asses when local sections of topological polymers (such as long DNA chains) might be able to detect global knotting of the structure (finding often local-writhe metrics can do so).

Writhe-based measures for proteins have been proposed although their use is less developed in this field as it is for DNA. The first example of which we are aware is in [54, 55], where the writhe and its generalisations are used to effectively classify proteins in agreement with the CATH database to a high degree of accuracy. The generalisations are the so called higher order Gauss integrals. If the integrand density of 1 is written as *w*(*s, t*), then the higher order variants are in the form:

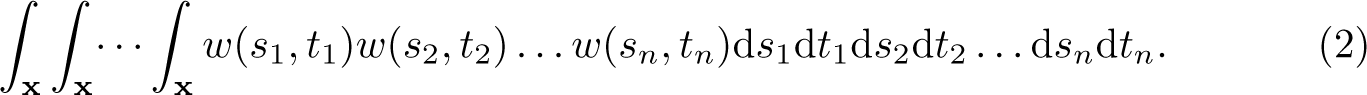

Unlike the standard writhe measure these quantities can be somewhat difficult to characterise. Intuitively it can be thought of as the number of times the directions of labelled pairs of crossings, say (*s*_1_*, t*_1_) (*s*_2_*, t*_2_), coincide (see [55] for a more detailed discussion). The authors use up to 30 of these variants to create a structural distance metric for comparing two structures, it is this metric which is shown to predict the CATH classification well in [54]. In [56] a comparison metric based on writhe value of subsets of the proteins of fixed lengths were combined with a number of other metrics to create another comparison metric against the SCOP classification, again with good results. In [57] a more rapid metric was created from the same Gaussian integrals as in the other two studies, and it was once again shown to compare favourably to the SCOP benchmark set. We highlight the fact that the difficulty in interpreting these higher order writhe quantities intuitively is one reason we seek alternative writhe metrics (despite their evident success). Further, one of our aims is to develop methods which can identify structures which are **not classed as similar using standard classification methods**.

The measures introduced in [54, 55] were recently developed further in [58] where the Gaussian-integral approach was applied to subsets (fragments) of the protein, via a fingerprinting technique, using the entanglement of sub chains to identify rare conformations in proteins, essentially pairs of subsections of protein which are mutually entangled in some manner which is relatively rare. By contrast, in our case we aim to demonstrate there are tertiary structure motifs which are both common and similar across proteins, but which are not classed as similar in the standard classifications.

One final aspects of the writhe literature we focus on in this is study is whether there are limits to the amount of entanglement in proteins as measured by the writhe. A theoretical upper bound on the writhe of smooth (thick) knots is presented in [59]:

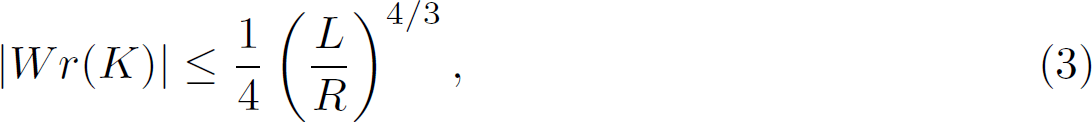

where *L* is the knot *K*’s length and *R* its radial thickness (so thicker knots are harder to entangle). Given proteins have distinct end points and are discrete curves, this bound does not strictly apply to proteins, in addition the notion of radius is a difficult notion to define for proteins (although some attempts have been made in the past [60]). In [61] the relationship between the writhe and length of uniform random walks is studied, and then applied to proteins in [62]. Since the local geometry of the protein’s backbone is tightly constrained by the Ramachandran steric constraints [63], we may expect that a random walk cannot capture the systematic entanglement of a protein’s backbone. In addition a very important characteristic of proteins is that many of the secondary structures such as *α* and strand helices are highly rigid, which significantly constrain their potential for creating complex global folding [64]. This motivates one of the novel aspects of our writhe metric, that we smooth/coarse grain the backbone model by the number of secondary sections of the protein, thus treating these secondary structures as rigid edges. We demonstrate using this coarse-graining that there are clear empirical constraints on the folding of protein structures as a function of the number of secondary structural elements of the backbone. Further we show our metric can be used to identify when secondary structure classification in a PDB is suspect.

As a final comment on the use of the writhe to characterise protein structure, we feel some of the properties of the quantity have been so far somewhat ignored in the protein literature. Firstly the writhe of helical geometries grows linearly with length, we will use this factor to identify and compare super-structure scale helical geometries such as those shown in Figure 2(c) and (d). Second, the writhe is a signed quantity, it can be zero for highly entangled curves. In [57] low or zero writhe measures were explicitly ignored from the comparison metrics, we show here, however, there are a number of interesting repeated zero writhe geometries which arise in protein tertiary structures which mimic techniques used to prevent cable entanglement and aid the creation of surgical sutures under confinement.

### Aims

To round up this somewhat lengthy introduction we detail the properties of the metrics we seek to develop and why they are novel to the field.

1. The metric should compare the similarity of tertiary structures ignoring their local secondary structure geometry.
2. Two structures should be classed as similar if they can be distorted into each other without significantly changing the topology of the fold (*e.g.* the preservation of helical geometry or knotting type), but allow for reasonably significant TM-scores between the structures.
3. The metric should pick up potentially meaningful structural similarities missed by the standard classifications.
4. The metric should provide clear bounds on the amount of entanglement exhibited by protein structures. This is necessary to ensure tertiary structure searches for relatively low-resolution experimental techniques such as BioSAXS can be restricted to plausible structures

In the methods section we discuss the two main quantities the SWRITHE package calculates, the writhe and absolute crossing number (unsigned writhe). In addition we discuss the properties of these two quantities and how they relate to specific features like superhelical geometries and zero writhe structures. Finally we discuss the motivation and method for our coarse-graining/downsampling of the C*^α^* backbone and the procedure to do this whilst preserving the fundamental topology of the backbone. In the results section we fist show both quantities have highly restrictive empirical bounds across a large (*>* 10000) sample of structures (which differ significantly at the fingerprint level). We show how helical super-structures and knotting are the main ways of developing systematic complexity in structures. We show how the unsigned writhe bound can be used to restrict expansive structural search methods for BioSAXS data.

We demonstrate there is a very common super-helical scale to tertiary protein structures which is common across a variety of CATH structure types, and that the example similarity of the TIM barrel and Rossmann fold discussed in this introduction is indicative of a consistent similarity between these two fold types. Finally we demonstrate there is a consistent zero writhe (but complex) structure across a variety of structure types which we believe merits further study.

## Methods

First we introduce the basic quantities calculated by the SWRITHE python notebook, then discuss some of their relevant properties. One can compute these quantities for a given protein using the accompanying code.

### The writhe

We consider a discrete curve *C* of length *j*, characterised by a set of three dimensional coordinates 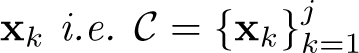. We calculate the writhe of a subset (sub fragment) of the curve 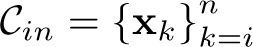 through the following formula:

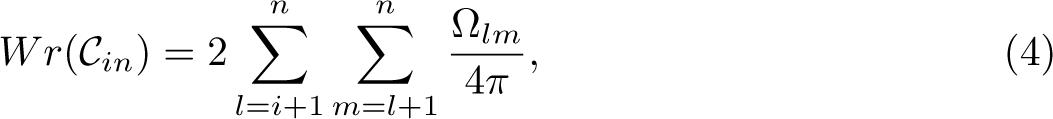

where Ω*_lm_* is a signed spherical area representing the contribution to (1) from crossings of edges connecting the pairs of points (*l –* 1*, l*) and (*m, m –* 1) (there are various formulae for Ω, we use method 1a of [65]). The idea behind the calculation of spherical areas is that crossings can be represented as points on the unit sphere and the writhe represents a signed area covered on this sphere (it can include multiple full coverings of the sphere), so the areas represent signed crossing densities [65]. The routine `calculate_writhe` calculates *Wr*(*C_in_*) for all 1 *≤ i ≤ j −* 5*, i* + 5 *≤ n ≤ j* for a given curve *C* (we need at least 5 points for a meaningful writhe calculation).

### The average crossing number

For bounds on entanglement in particular it is useful to count just the number of crossings as a measure of complexity of the fold. We denote this quantity as *acn*(*_ij_*) (the average crossing number):

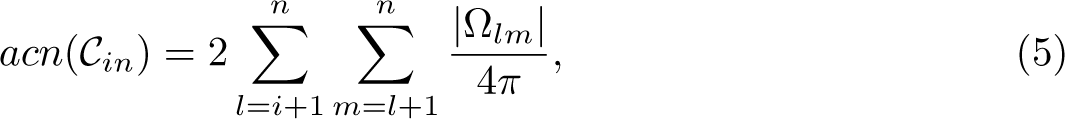

This quantity is also invariant under rotations and translations so needs no alignment. The routine `calculate_abs_writhe` in the SWRITHE notebook calculates *acn*(*C_in_*) for all 1 *≤ i ≤ j −* 5*, i* + 5 *≤ n ≤ j* for a given curve *C*.

### Aspects of the writhe critical to our study

#### Helical structures and linear writhe growth

Helices have a uniform writhe density (they have consistent chiral-coiling). That is to say *Wr*(_1*n*_), plotted as a function of subsection length *n* would be a straight line whose gradient can be given in terms of the helix’s pitch *P* and radius *R*. The writhe per turn is:

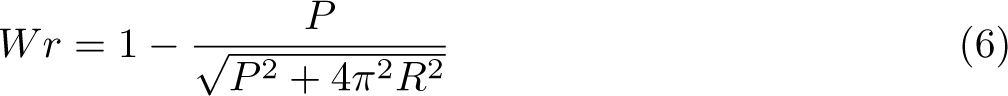

[67]. In Figure 5(a) we see an example of a smoothed C*^α^* backbone, smoothed again by only sampling one in every 3 amino acids, which has a clearly repeated helical geometry. In Figure 5(b) we see a plot of *Wr*(_1*n*_), the writhe of the smoothed C*^α^* curve as a function of its length. There is a clear almost linear rise due to the helical nature of the structure. Indeed, this structure has radius 7.58, and pitch 5.98 (found be approximating the helical axis), giving a writhe per turn of 0.88. Considering the substructure relating to the linear portion of the graph (that is between *n* = 30 and *n* = 130) there is a rise in writhe of 12. We compare this to the per turn result by looking at the actual structure, for which we count 14 clear turns. Since 14 0.88 = 12.3 it is clear this writhe calculation is accurately quantifying the consistent helical superstructure of the protein. We shall see later that this geometry is common to sub units of a large number of proteins, spanning many CATH domains.

**Fig 5.**
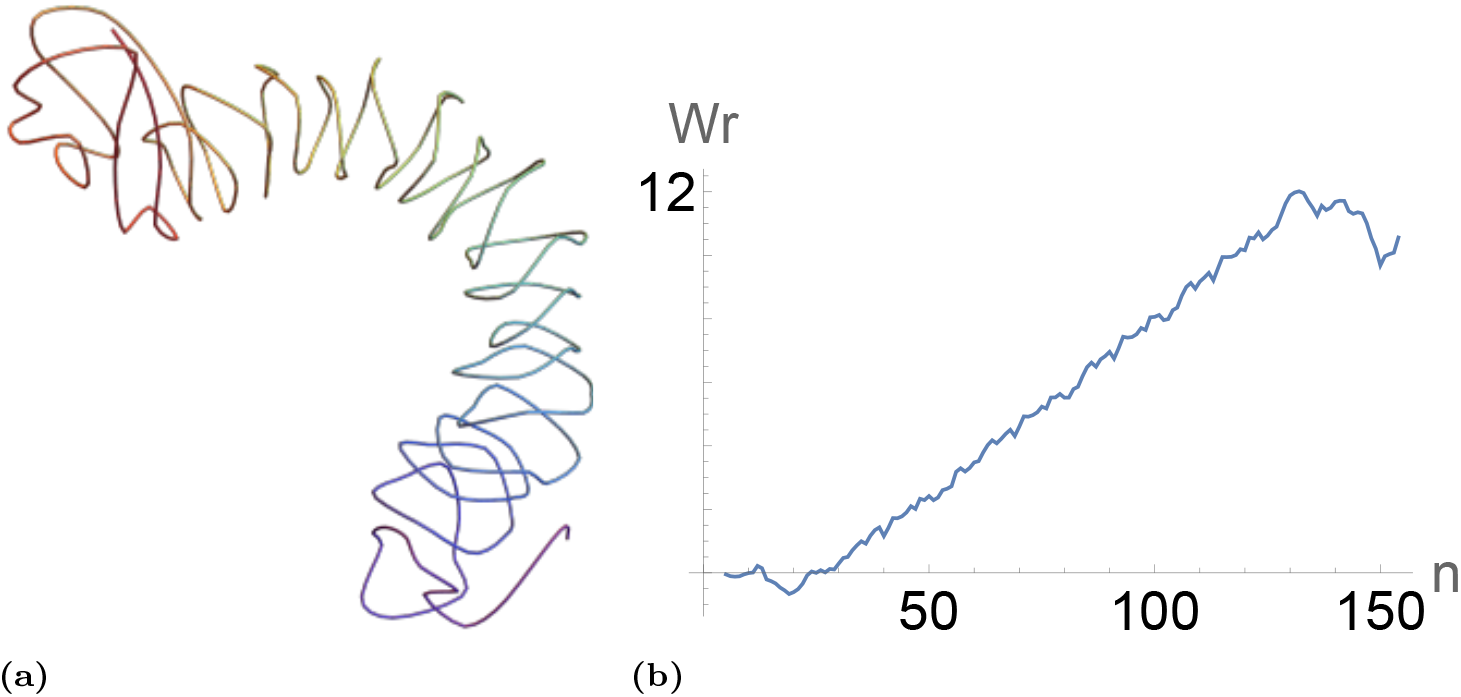
Helical writhe growth in a protein. Panel (a), the smoothed backbone structure of the Y369A/hEC1 complex (PDB 2OMZ) [66]. Panel (b) the writhe *Wr*(*_in_*) of the smoothed protein (selecting every 3 amino acids).

#### Secondary coiling

A word on the helical nature of secondary structures like strand helices and *α*-helices is appropriate at this juncture. They naturally have significant writhe, but this is not of interest to us as we are trying to classify the writhe/knotting similarity of tertiary structures. This is why the curve in the previous example was sampled every three amino acids. In Figure 6 we see a structure which is dominantly *α*-helical, its writhe as a function of length is charted in panel (b) and is nearly linear and positive, reflecting the positive chirality of the *α*-helices. The smoothed structure, sampled every three amino acidsm is shown in (c). A rough left-handed helical coiled can (with some effort) be identified, this is reflected in the roughly linear build of up negative writhe seen in (d). Sampling every 4, 5, 6 & leads to a similar plot as that shown in panel (d) indicating this (distorted) left handed coiling is a persistent signature of the tertiary folding of this structure. This intricacy has been noted before, for example in [57], where a centre-of-gravity type smoothing is proposed in order to reduce the contributions of small scale helices to the writhe signal. While this method is effective, for our purposes we will propose an even coarser smoothing which puts greater emphasis on the large scale entanglement of secondary structures.

**Fig 6.**
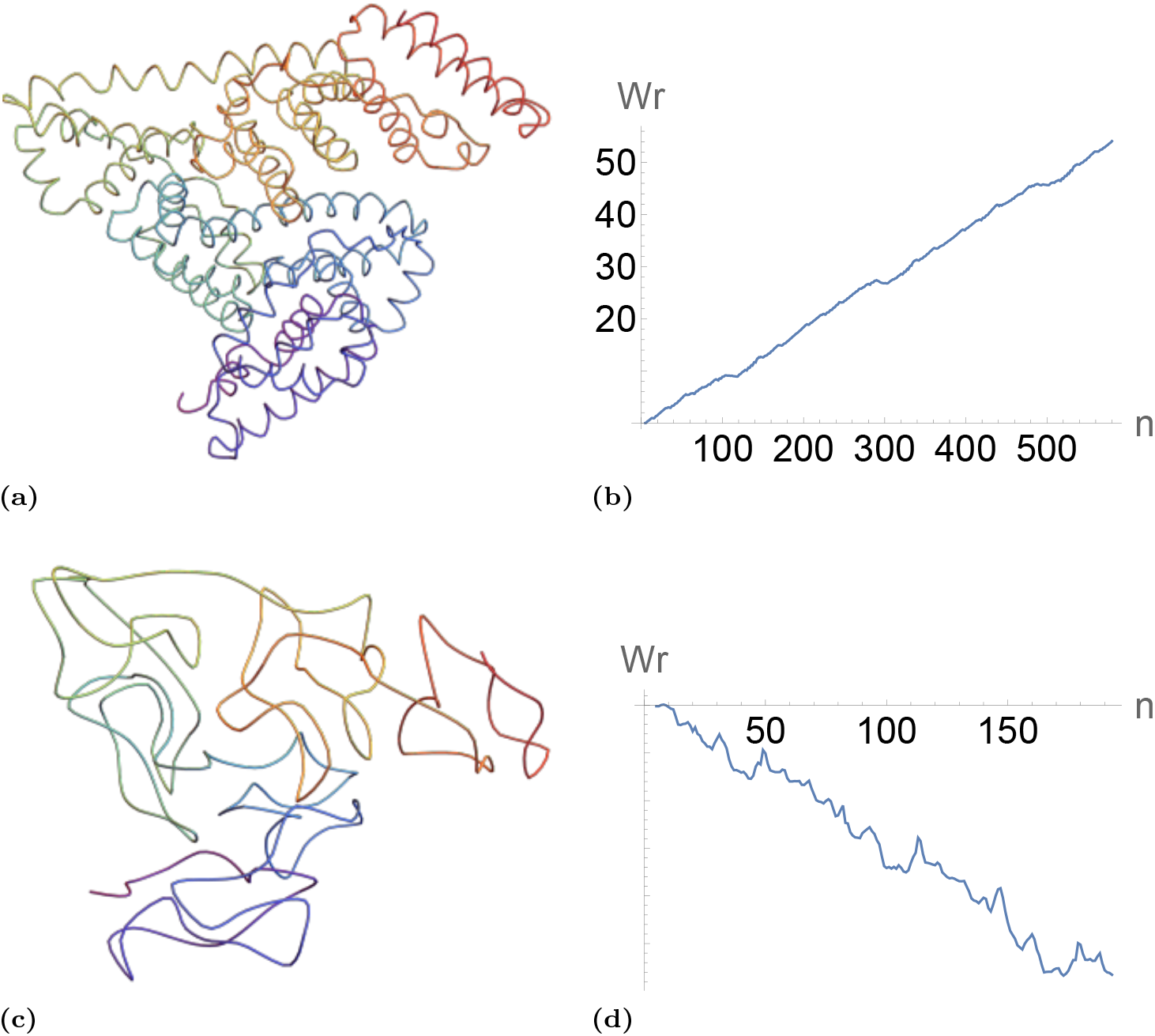
Illustrations of the effect of smoothing the backbone to avoid secondary structure writhe. Panel (a), the C*^α^* backbone of Bovine serum albumin (pdb 3V03 [68]), it is a dominantly *α*-helical structure. Panel (b) the writhe as a function of length for the structure in (a) it is nearly liner with a build up of significant positive writhe, commensurate with the *α*-helical geometry. Panel (c) is the curve in (a) sampled every three amino acids, the secondary structure is smoothed out, its hard to see but there is a large scale helical coiling present. Panel (d) is the writhe as a function of length. It is roughly linear commensurate with the larger scale helical coiling.

#### Meaningful zero writhe and “roadie” geometry

It is possible for a structure to have close to zero writhe whilst still being very entangled. A fact that is very familiar to sound engineers, who have developed a method of storing cables which ensures they can be easily uncoiled by simply pulling at the coils end’s. A cartoon depiction of this method is indicated in Figure 7(a), where the combination of over/under windings yield crossings with alternating sign, meaning the net entanglement is zero. An idealisation of this geometry is shown in 7(b), we see in (c) its writhe *Wr*(_1*n*_) rises and then falls as a function of *n* leading to zero net writhing. This geometry is also a geometry used in slipknot sutures as it is easy to slide along the thread and create knots in tight spaces [71]. So the writhe measure is correctly quantifying the ease to which the structure it can be disentangled. This motif is observable in proteins too, in 7(d) we see a highlighted section of a protein whose backbone has the over-under coiling structure required to have close to zero writhe. The average crossing number *acn* by contrast would yield a positive value for this geometry indicating it is non-trivially tangled.

**Fig 7.**
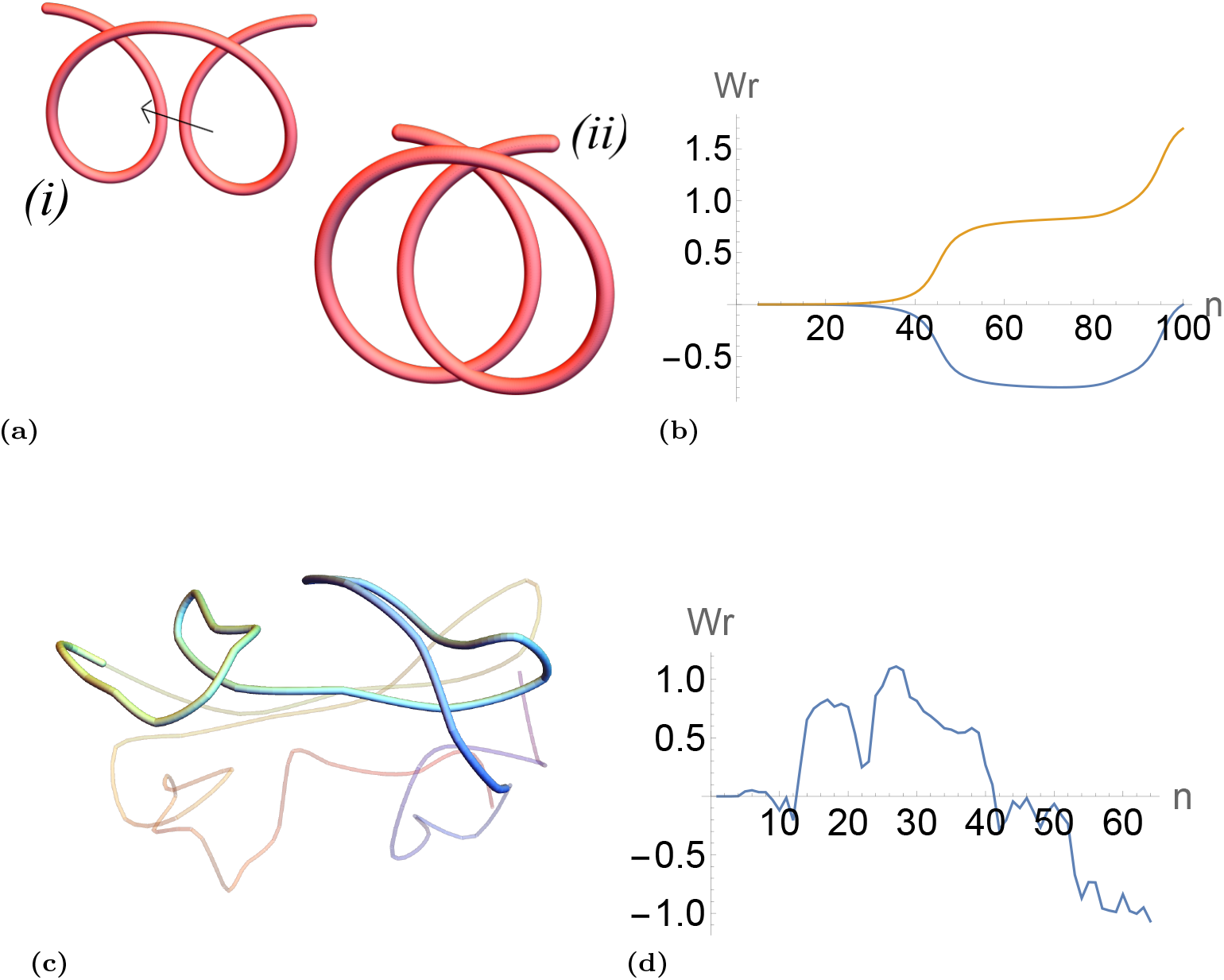
The “Roadie wrap” geometry, which illustrates the utility of the signed (orientational) nature of the writhe [69]. Panel (a) is a depiction of the method for storing cables which ensures they can be easily disentangled, as indicated subsequent loops cancel in their writhe contribution Two counter helical loops (i) are created, the bought together in (ii) (the arrow indicates the second loop is bought in between the first. Panel (b) which charts its writhe *Wr*(*C*_1*n*_) of curve (i) in (a), and its absolute crossing number *acn*(_1*n*_)), as function of length. The writhe plot which increases then decreases back to zero whilst the crossing number ends up with significant entanglement. Panel (c) is a smoothed backbone of the Human Inosine Triphosphate Pyrophosphatase (PDB 2I5D) [70]). The section highlighted has the same geometry (in reverse) as panel (a). This is clear in its writhe plot (d) where the section between *n* = 12 and *n* = 38 leads to no overall writhe but has significant values half way in between.

In this study we are unable to offer a explicit functional reason for this particular geometry to exist in proteins, unlike for the strictly helical geometry, but we show instead that it is highly prevalent in numerous PDB structures. We use the following fairly strict criteria for locating a roadie-like geometry. We require a section *Wr*(*C_in_*) for which *Wr*(*C_in_*) *<* 0.05 and for some *Wr*(*C_im_*)*, m < n* we have *Wr*(*C_im_*) *>* 0.90. The second criterion ensures that the section *_in_* has significantly coiled/helical topology which is subsequently cancelled, thereby ruling out sections with little complexity. We note this would include the example in 7(c) and we seek similar geometries. Later in the results section we find there are a significant proportion of proteins for which this is true. In what follows, when we refer to a “roadie” geometry we are explicitly referring to subsections *C_ij_* which meet these criteria.

#### Integer changes and crossing detection

The writhe changes by an integer value when the curve crosses itself (±2), thus it can be used to detect crossings when curves change shape. This property makes it a critical quantity in DNA topology studies where it can be used to identify and ignore non physical deformation path of DNA models [44, 45, 72]. This property ensures that the writhe of the figure 8 curves shown in 3 are respectively +1 and 1, so the structures are classed as fundamentally different, by contrast to the RMSD metric. Another example of the utility of this property is indicated in Figures 8 and 9. Figures 8 depict snapshots of a continuous deformation of a circle into a trefoil, a path which must necessarily include self intersection of the curve. In panel 9(a) we see the writhe of the curve plotted as a function of the deformation (parameterised from *t* = 1, the circle, to *t* = 101, the trefoil. We see numerous sharp jumps which as shown in panels (c) and (d) correspond to self intersections, by contrast in (b) we see the RMSD distance between neighbouring configurations *t* and *t* + 1 changes smoothly and does not characterise these sharp changes in entanglement (it is normalised by the mean distance between points on the curves).

**Fig 8.**
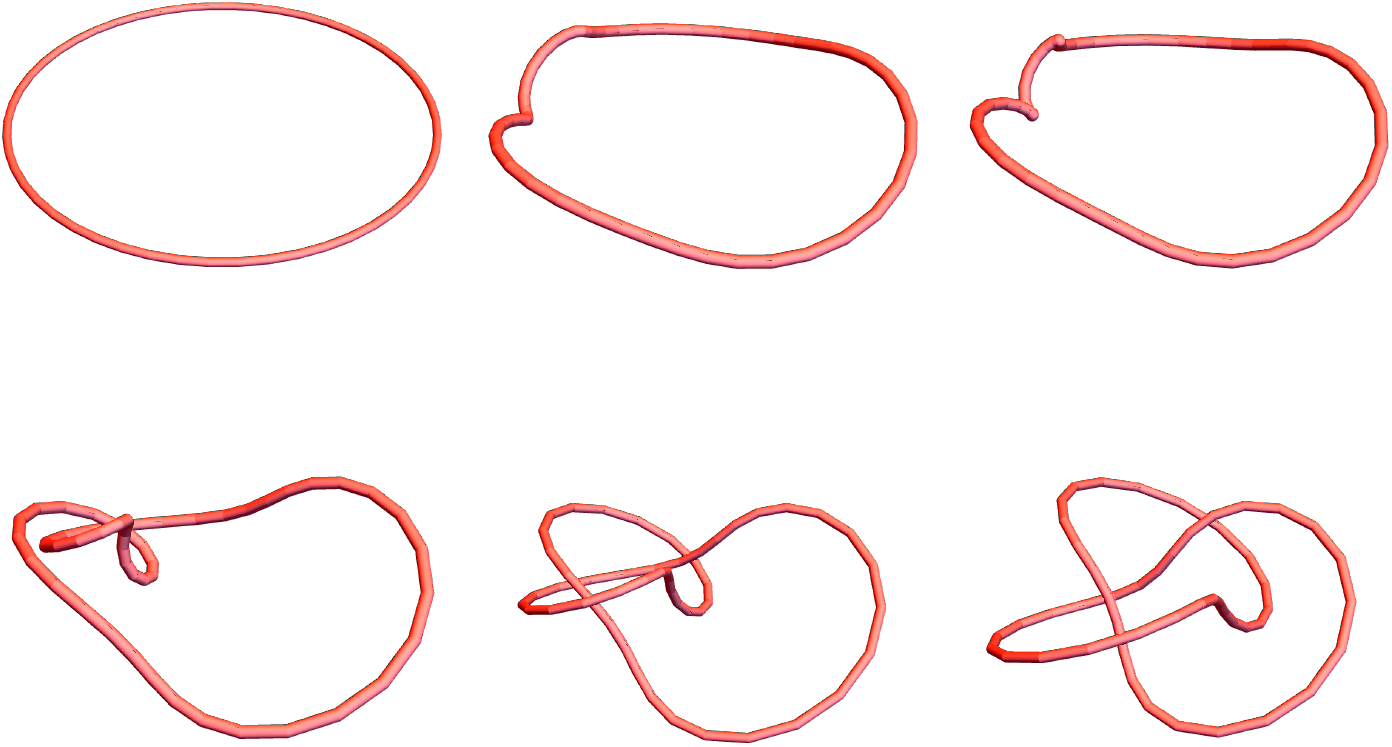
Selected configurations of a continuous deformation of a circle into a trefoil, shown ordered from left to right, top to bottom. The curves are discrete, composed of 50 edges. There would necessarily be some points during this deformation path where the curve crosses itself.

**Fig 9.**
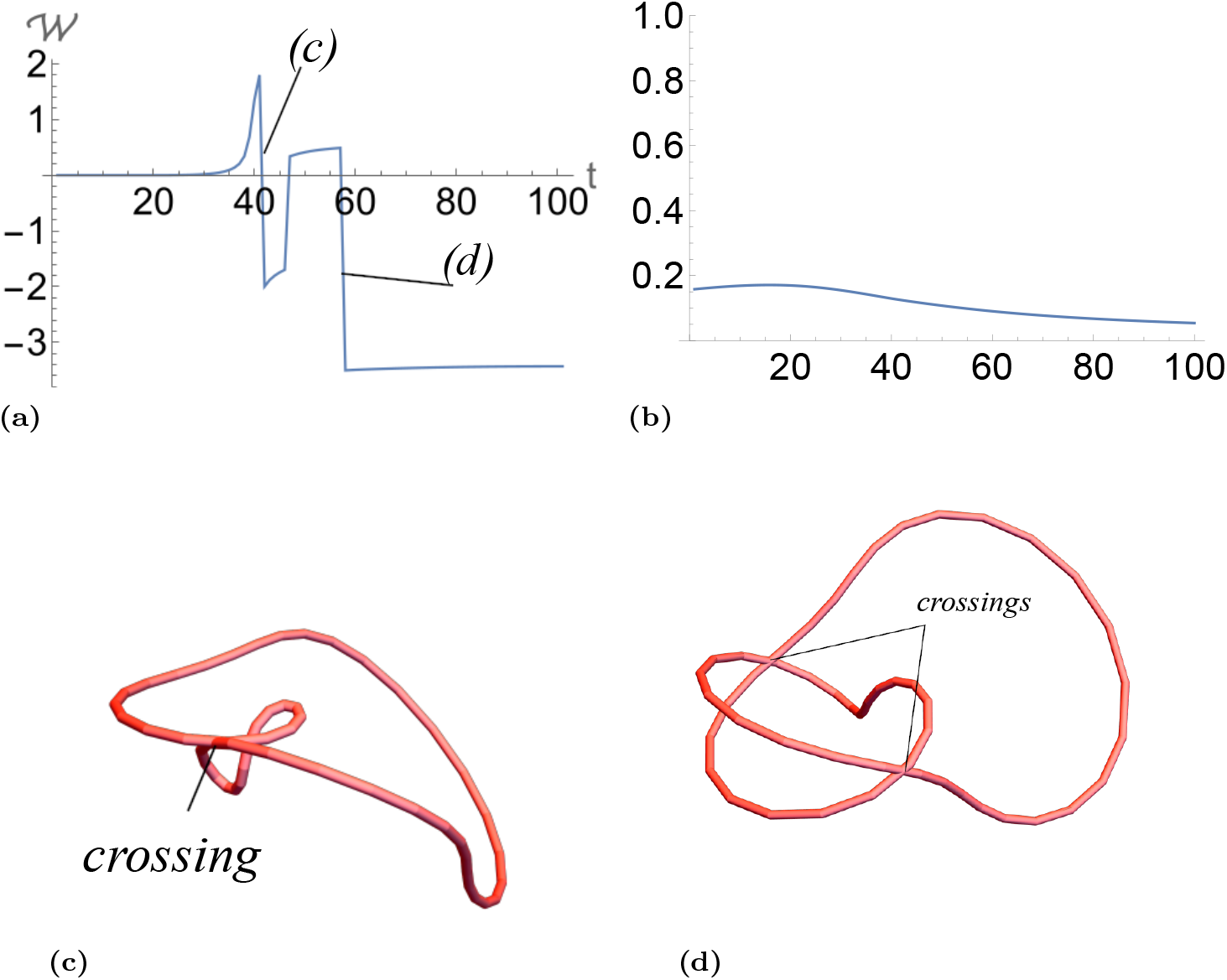
Evaluations of the deformation shown in Figure 8. Panel (a) is a plot of the writhe as a function of *t* parameterising the deformation. Two of the discontinuous writhe jumps are labelled which correspond to the crossings shown in panels (c) and (d). Panel (b) is the RMSD between neighbouring configurations (normalised by the mean segment distance of the curves) it is continuous.

We can see this property’s utility by considering the homologous pair of proteins detailed in [73], 3KZK which forms a trefoil knot in its core, and 4JQO which is unknotted. These proteins are examples of AOTCase and OTCase respectively. As shown in [73], these two proteins are structurally very similar, differing only by a crossing change. Since these two proteins can be easily superimposed, the RMSD between them is small at just 1.71Å. That is to say, this difference isn’t seen as a significant. In 10 we plot the writhe of these two proteins as a function of length. One can clearly pick out where this crossing change occurs. Similarly to the trefoil example above, the crossing change leads to a significant jump in writhe, as the two states are fundamentally different. As noted in [74], this structural change is biologically important, changing the size of the binding pocket. This difference is picked up via the persistent homology based metric in [73], highlighting the fact that the writhe can detect these important topological features that more sophisticated (hard to compute) measures identify.

We also intend to show the counter property, that the writhe can be used to detect a similarity of structures that it is not apparent via RMSD based measures. But this is a more subjective notion and will require more context and evidence to argue.

### Smoothing by secondary structure

#### The rationale

We are aiming for a measure of the writhe of protein backbones which does not include the secondary structure’s inherent helical nature. One might suggest we sample the full C*^α^* backbone curve of the protein every 3 or 4 amino acids as in the previous examples shown, this is the procedure used in [57]. However, we argue a better choice is to represent each secondary structure element (SSE) as an edge of a discrete curve. For *α*-helices and *β*-strands this argument is clear a they are reliably inflexible structures whose contribution to the overall tertiary fold topology/geometry can only be as an inflexible edge.

For these structures we can safely represent each SSE by a single point at its first residue and the next point on the next secondary section to which it is joined. We would like to argue the same for the more varied linker geometries also. Often, whilst they have more variable local geometry their main functionality is to link secondary sections (as in *β* sheets) and the specific geometry of the section is less important than the orientation of its end points in joining the helical secondary structures. However, there are occasionally linker sections which coil around other SSEs leading to knotting (or slip knotting and the other various rare but complex entanglements found in some proteins [75]). So replacing them by a straight edge will miss this essential entanglement. We have created an algorithm for identifying the minimum number of points in a linker SSE which preserves any such knotting with a minimal number of points, as in Fig 11.

**Fig 10.**
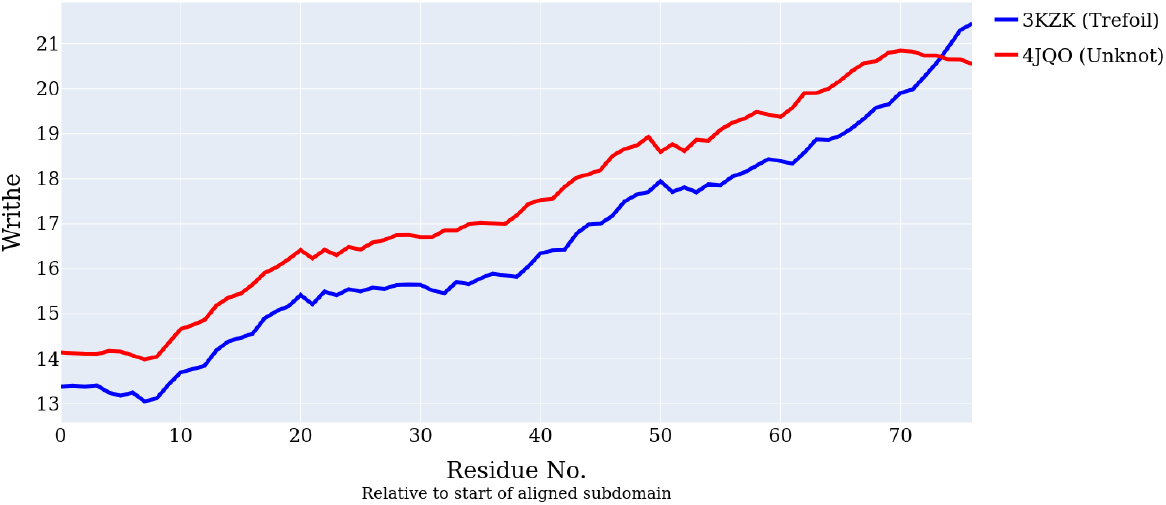
A plot of the writhe for the aligned subdomains of 3KZK and 4JQO. The crossing change between residues 70 and 80 leads to a different topology between the two structures, and is identified by the writhe.

**Fig 11.**
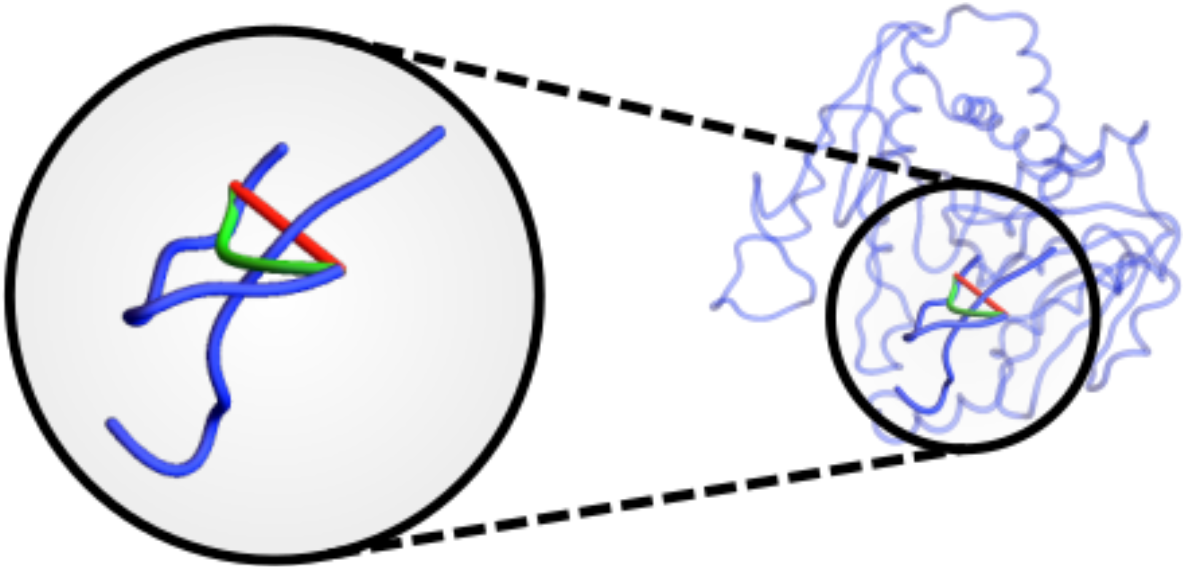
An example of the non local coiling that is preserved via the SKMT algorithm. In blue we see the backbone of the trefoil knotted acetylornithine transcarbamylase complexed with acetylcitrulline (PDB 3KZK) [76]. In red we highlight the straight edge connecting consecutive SSEs, which passes under the C terminus. In green, we see the edge connecting these SSEs output by the SKMT algorithm, which preserves the non local crossing, and therefore the knottedness of this curve.

#### The method

We construct a new discrete curve from the full C*^α^* backbone as follows. We assume as input the coordinates of the C*^α^* backbone and a secondary structure assignment.

1. Take the coordinates of the N-terminal amino acid.
2. For the next secondary structure element, if it is an *α*-helix or *β*-strand we select its first point
3. If it is a linker we find the minimum number of points required to preserve any knotting it has with the rest of the curve. To do so we use a method similar to the KMT algorithm [40], if the triangle defined by three sequential points is not intersected by any edge of the curve, then we can safely remove the middle point of the triangle. See Fig 11 for an example of such a triangle intersection.

We call this the SKMT -secondary KMT algorithm in what follows. In Fig11 we see an example of this smoothing method for a subsection of the trefoil knotted 3KZK.

The SKMT algorithm can be applied to a selected PDB file using the skmt routine in the accompanying SWRITHE notebook.

#### Secondary section number as a length

As noted in [64], smaller proteins have very little scope to become entangled, since their SSEs are relatively large compared to the length of the protein itself. For a larger proteins, the relative length of its rigid subsections is often (but not always) smaller, and we can therefore see greater global entanglement. It makes sense then to consider entanglement scaling as function not of length/number of residues, but as a function of the number of SSEs. If we were to smooth a backbone simply by sampling every *k* points, at some points we would jump across SSEs, and therefore not accurately capture the entanglement as it relates to the arrangement of the rigid SSEs. We use this measure of length: the number of secondary structure sections, to derive empirical constraints on the complexity of proteins in the results section.

## Results

### Length constraints on the writhe of proteins

To study the distribution of writhe amongst proteins, we take a representative sample of proteins from the PDB via the following criteria:

1. A good resolution (*<*2Å)
2. Consisting of between 30-300 residues.
3. Redundancies removed at 70% sequence identity
4. Good model quality: *R_work_ ∈* [0, 0.2] and *R_free_ ∈* [0, 0.25].

This yielded 10736 entries from the PDB. We took the largest monomer unit from each PDB and obtain a simplified backbone using the SKMT algorithm described in. We too the “length” *L* of this curve to be simply the number of secondary structure elements.

The values of *W(C)*, the writhe of the whole SKMT curve, are presented in Figure 12 as a function of their length *L*. There are a number of critical aspects of this plot we highlight.

**Fig 12.**
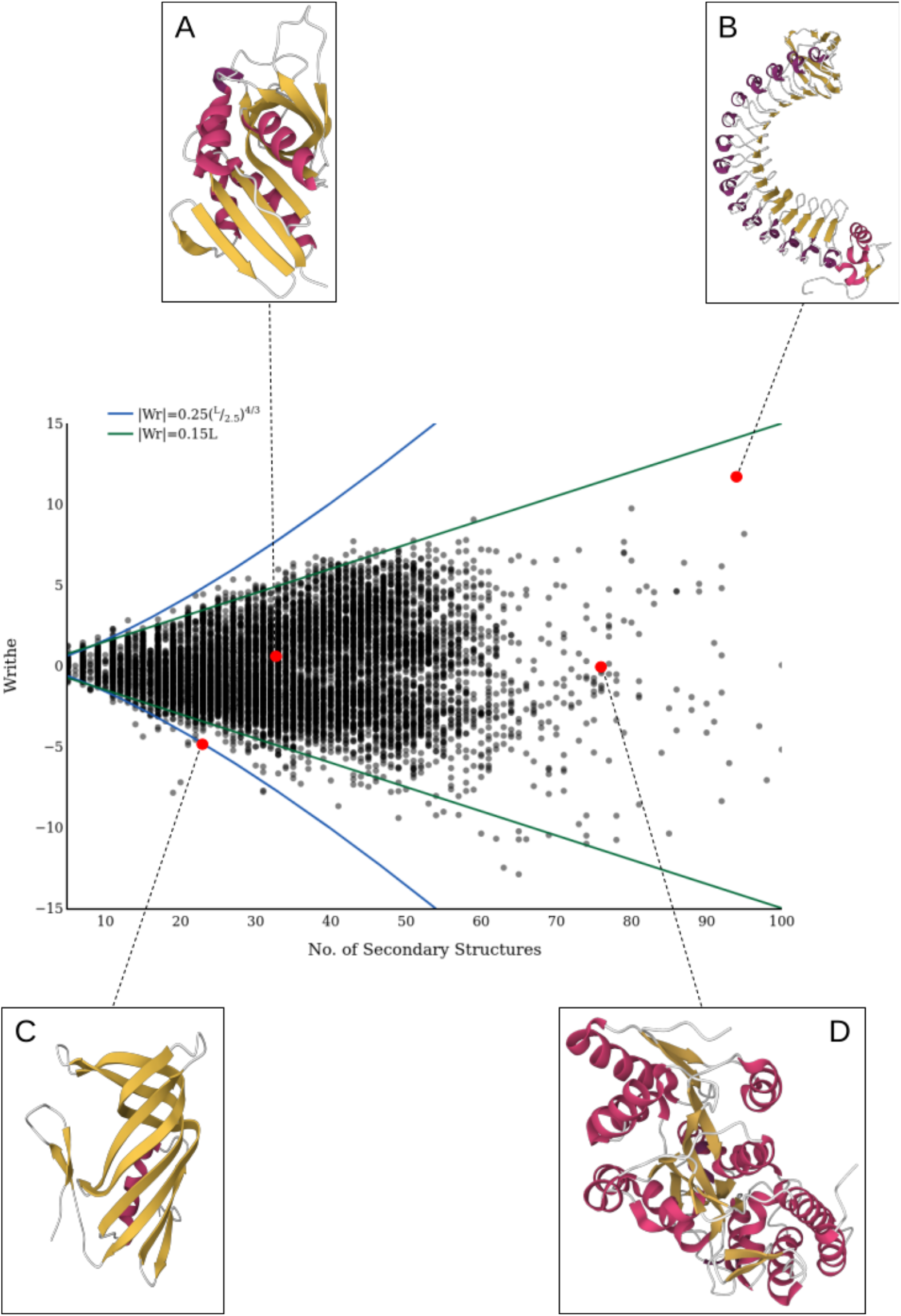
The distribution of writhe for a representative sample of 2298 proteins from the PDB in black. In blue we see the theoretical writhe bound [59] with *R* = 2.5. In green, we see linear growth in writhe with a gradient of 0.15. Inset: A: PDB entry 4QFB [77]. B: PDB entry 2OMZ [78]. C: PDB entry 3HWU [79]. D: PDB entry 4O4B [80].

**Fig 13.**
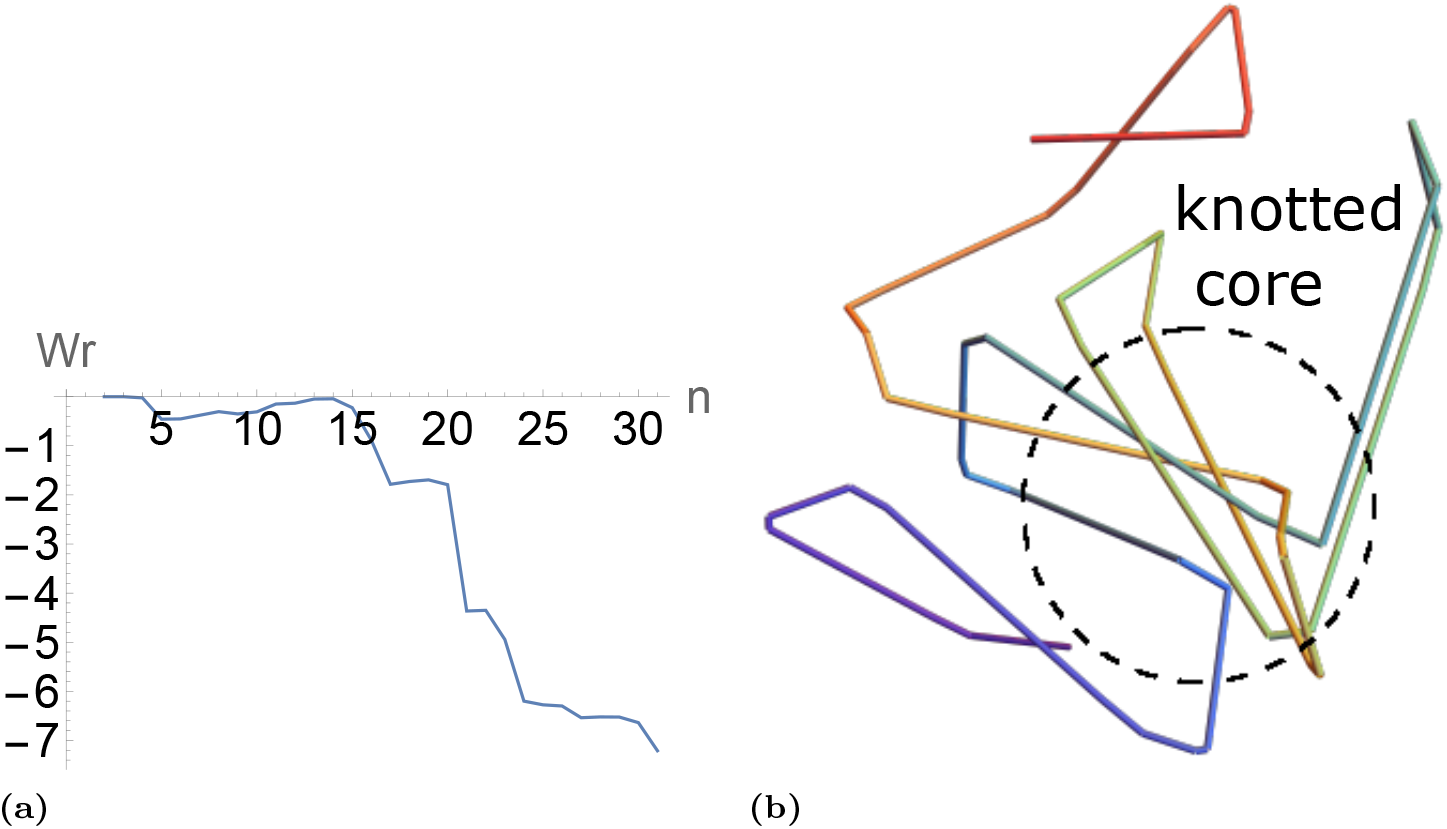
Knotting in a protein close to the knot bound. Panel (a) is reprsents *W* (_1*i*_) for [2*, n*] of the SKMT smoothed protein shown in Figure 12(C) which is close to the knotted bound, We see a heavy build up between sections 5 and 30. This can be seen in panel (b) to result from a knotted core of the protein (this protein was classified as a trefoil knot via the Knoto-ID software [81]).

1. 99.8% of all values lie within the curves:

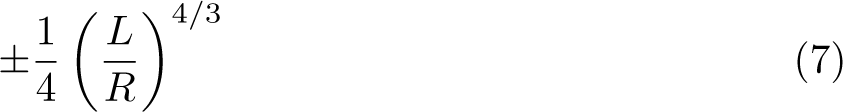 The theoretical knot bound from [59], with a radius of *R* = 2.5 (similar to a mean “tube” thickness of 2.7 Å found in [82] for minimal C*^α^* triplet radii).
2. For larger proteins, *L >* 35, the *Wr* values increasingly fail to get close to this limit. We find 97.4% of the structures fit within a linear bound 0.15*L*.
3. A significant number of proteins have a low or even close to zero writhe, consistent across all scales. An example is shown in Figure 12A.

The first knot “bound” assumes that the complexity available to a curve/tube increases with length, whilst a linear trend indicates a fixed complexity. independent of length, so it seems quite meaningful that such a high percentage fit inside this linear trend. Indeed. the helical protein shown in Figure 5, which is the same structure as shown in panel Figure 12B, can be seen to be very close to this linear bound. Note that the nearly zero writhe protein in panel A has many anti-parallel *β*-strands, with short linker sections between them, which leads to cancellation in the signed sum. In contrast, the *β*-strands of the protein seen in Fig 4B are all parallel, and separated by a linker-*α* helix-linker pattern As a result, there is a consistent handedness of winding along the length of the protein, leading to an almost linear build up in writhe. This protein is an example of a Leucine Rich Repeat (LRR) Right-handed Beta-Alpha superhelix, as per the CATH topology classification [7].

Fig 12C shows a much smaller protein whose writhe sees super-linear growth (it is on the “knot” bound). It is clear that the main contribution to the writhe for this protein is its *β*-barrel like structure. One can see in Fig 14 its writhe grows rapidly due to a knotted core in the second half of the structure. By contrast Fig 12D highlights that it can be quite difficult for a protein to build up systematic writhing. The structure is a large protein with a visually very complex entanglement.The total writhe of this protein however is close to 0. One can see in Fig 14(a) it has a peak writhe value of 2 which which arises from a locally helically coiled geometry at the beginning of the structure (Fig 14(b)). But it also has a zero writhe “roadie” type structure (Fig 14(c)) adding complexity but no net writhe, and then a long near linear section (Fig 14(d)) which passes through the middle of the structure cancelling the previous writhe. Such complexity is quite common for a large number of structures in this data set. We show later in how the SWRITHE code can be used to identify both helical and “roadie” substructures in a given protein structure.

**Fig 14.**
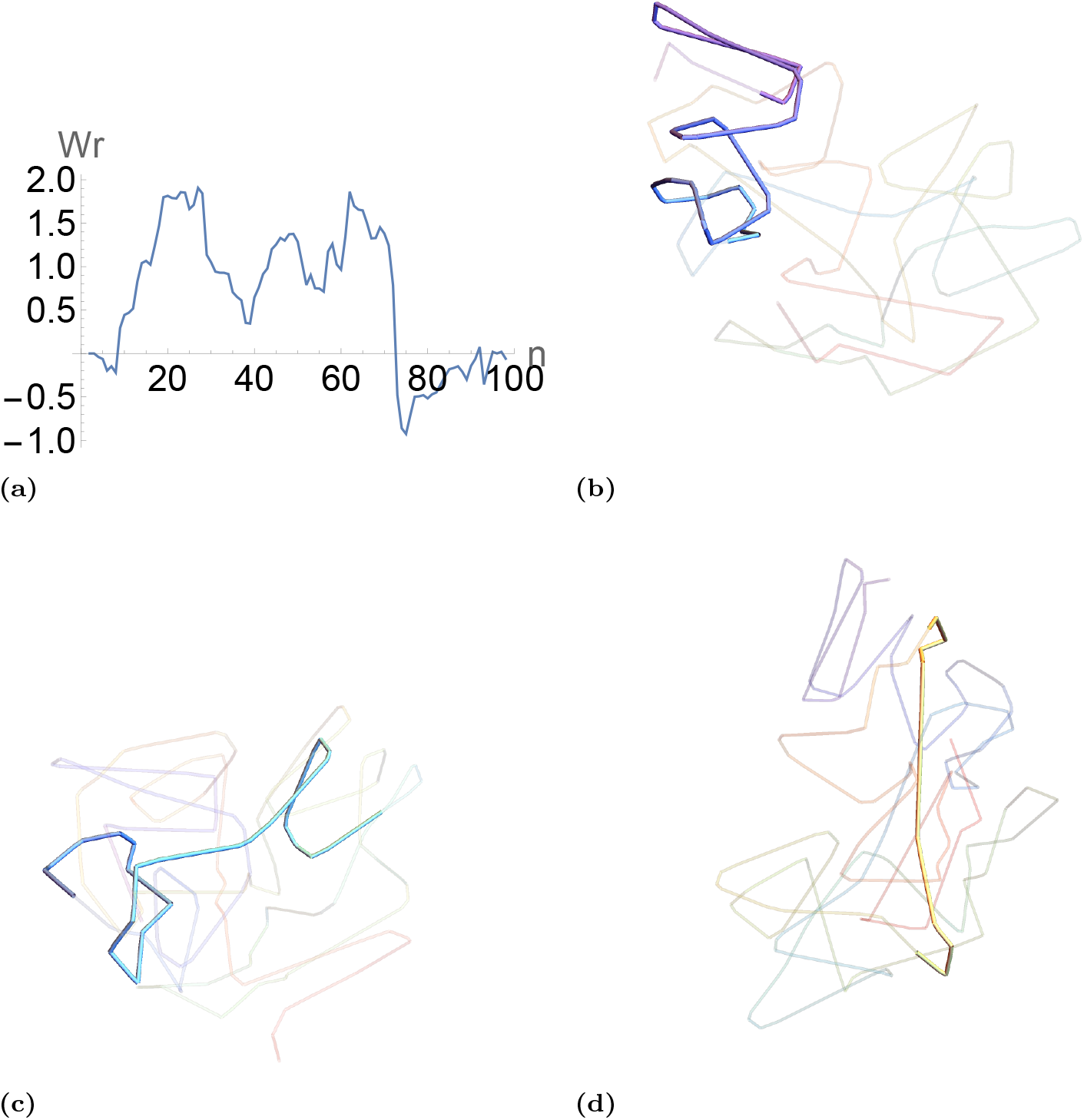
A complex (near) zero net writhe structure. Panel (a) shows the development of the sub-sectional writhe *W* (_1*i*_) for [2*, n*] of the SKMT smoothed protein shown in Figure 12(D). In panel (b) we see the structure with sections 19-24 highlighted. This corresponds to a roughly linear growth in writhe in plot (a) and can be seen to have a clear helical geometry with 2 turns (hence a maximum writhe of approx 2). Panel (c) shows a roadie like geometry in highlighted sections 18 − 37 this correspond to a growth and then decay of the writhe plot in panel (a) as one would expect for a (nearly) net zero (writhe) structure. In panel (d) we see highlighted a nearly linear geometry of sections 68-76 which corresponds to the sharp drop of almost 2 in the writhe plot shown in (a) as it passes though the middle of the structure (so is seen to add significant negative crossings).

### The distribution of writhe for trefoil knotted proteins

To further understand this distribution of writhe across the PDB, we look at a subset of knotted proteins. In particular, we consider the set of open-ended trefoil knotted proteins as detailed in [83]. In Fig 15 we see the distribution of writhe for this set of proteins (in red), compared to the distribution of writhe amongst our general data set (in black). We can see that for larger molecules *L >* 35 there is little correlation between trefoil knottedness and writhe. By contrast, for smaller molecules the presence of a knotted core yields a significant amount of writhing, particularly negative writhing corresponding to left handed trefoil knotted cores. In the case of the longer molecules the fact that a protein gets closer to its bound is often due to the presence of helical geometries, as seen in Fig 16. Since it is well understood that random walks are more likely to be knotted than proteins [35] these results make it clear that larger structures often generate systematic complexity through helical superstructures. In this way, we see that the writhe measure captures some information that is lost within the knot based measures.

**Fig 15.**
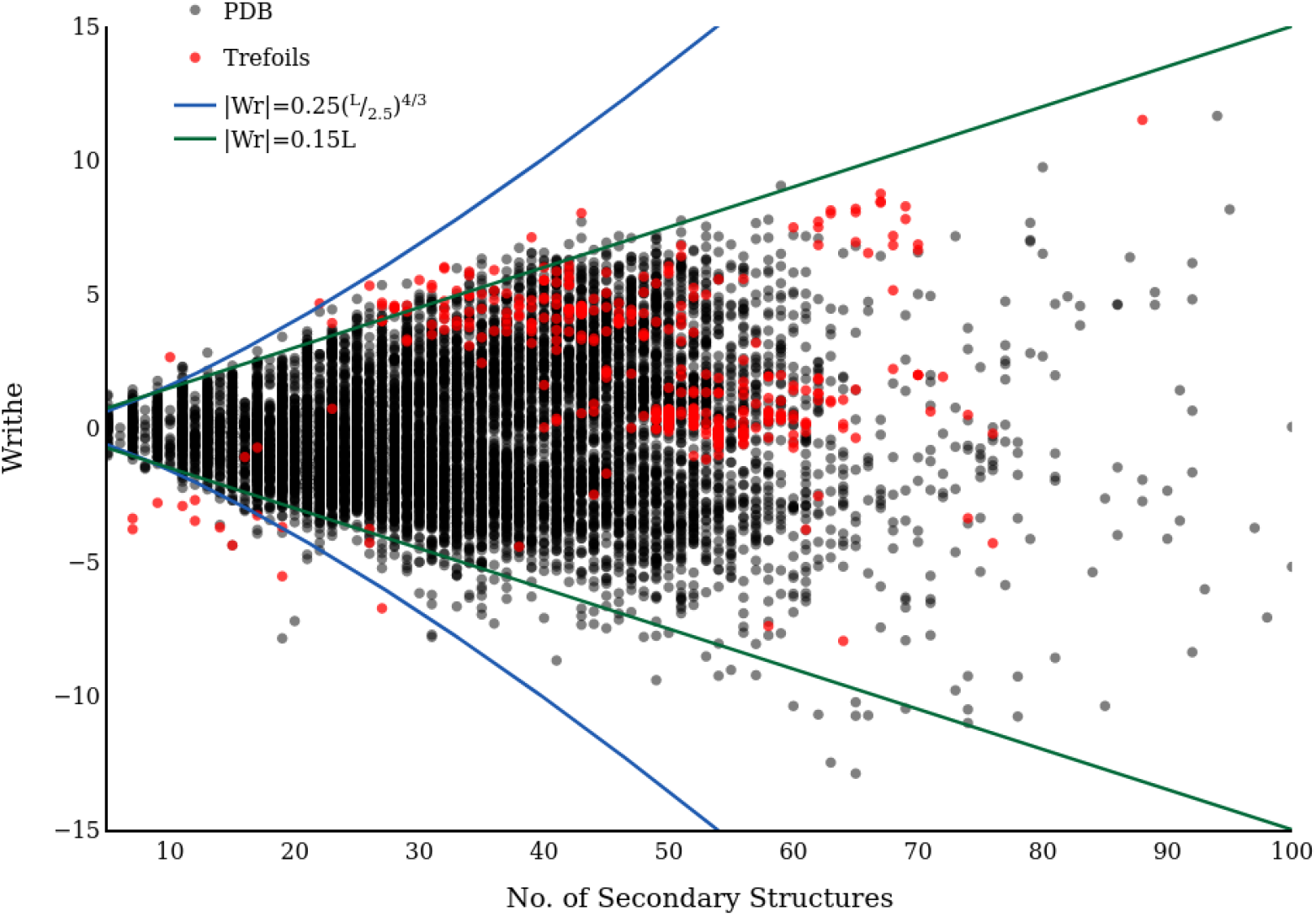
The distribution of writhe amongst the open trefoil knotted data set from [83] in red, compared to our subset of the PDB in black.

**Fig 16.**
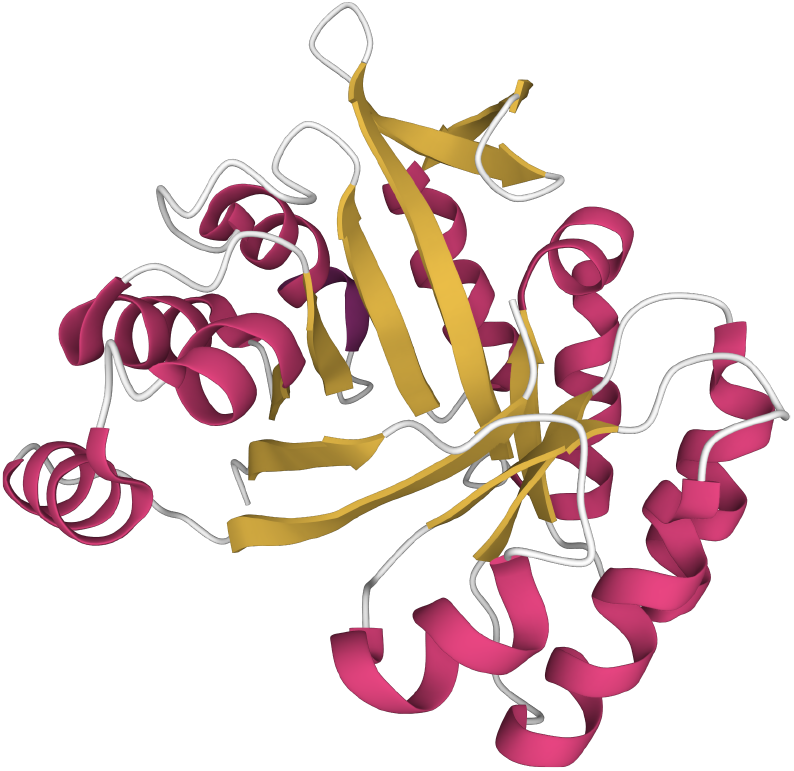
Cartoon representation of a beta/alpha-barrel built by the combination of fragments from different folds (PDB: 3CWO). This protein is an example of a trefoil knotted structure which has a globally helical structure, thereby maximising writhe.

### An empirical bound on the *acn* of proteins

As we have seen complex structures can have little or no net writhing, the average crossing number *acn*(*C*) then allows us to measure the net complexity irrespective of the sign. This means for example a double looped helix and the roadie type structure will have the same complexity measures, as indicated in Figure 17. In the context of determining structures using BioSAXS data, the *acn* is much better suited as a complexity penalty, as we will see below. We show now there are apparent empirical constraints on this measure of complexity for protein structures by calculating *acn*(*C*) for the same set of proteins as in the previous section. The distribution is shown in Figure 18

**Fig 17.**
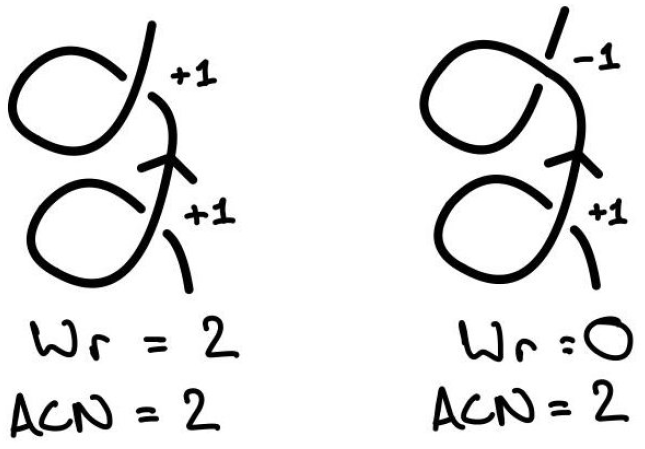
An example of the distinct properties seen by the writhe and *acn*.

**Fig 18.**
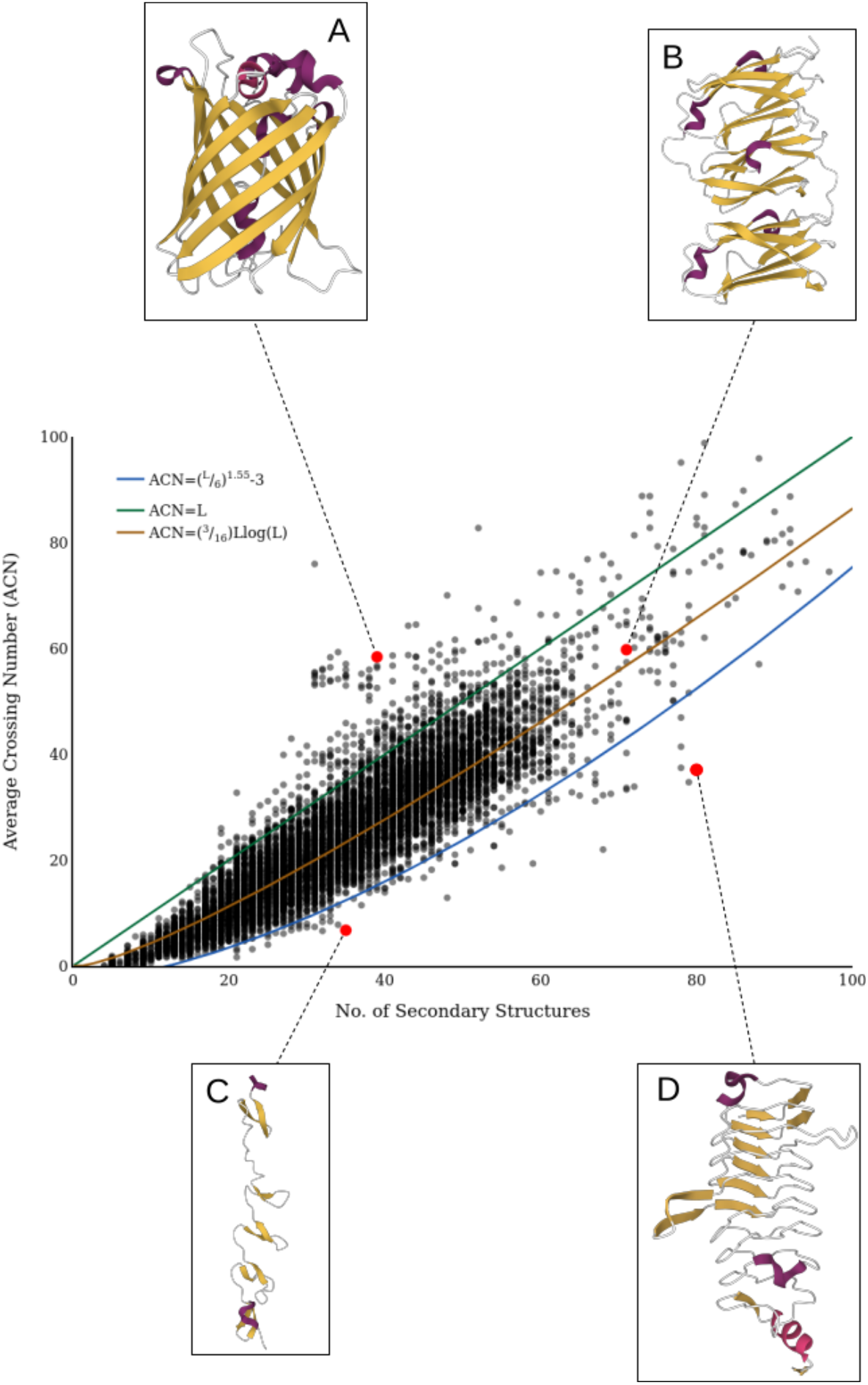
The distribution of *acn* for a representative sample of 10736 proteins from the PDB. In blue, the bounding curve described is the bounding curve. In orange, the *O*(*L* log *L*) growth in *acn* as in [84]. In green, linear growth in *acn* with respect to length. A: PDB entry 3EVP [85]. B: PDB entry 4M9P [86]. C: PDB entry 1V1H [87]. D: PDB entry 3PSS [88].

**Fig 19.**
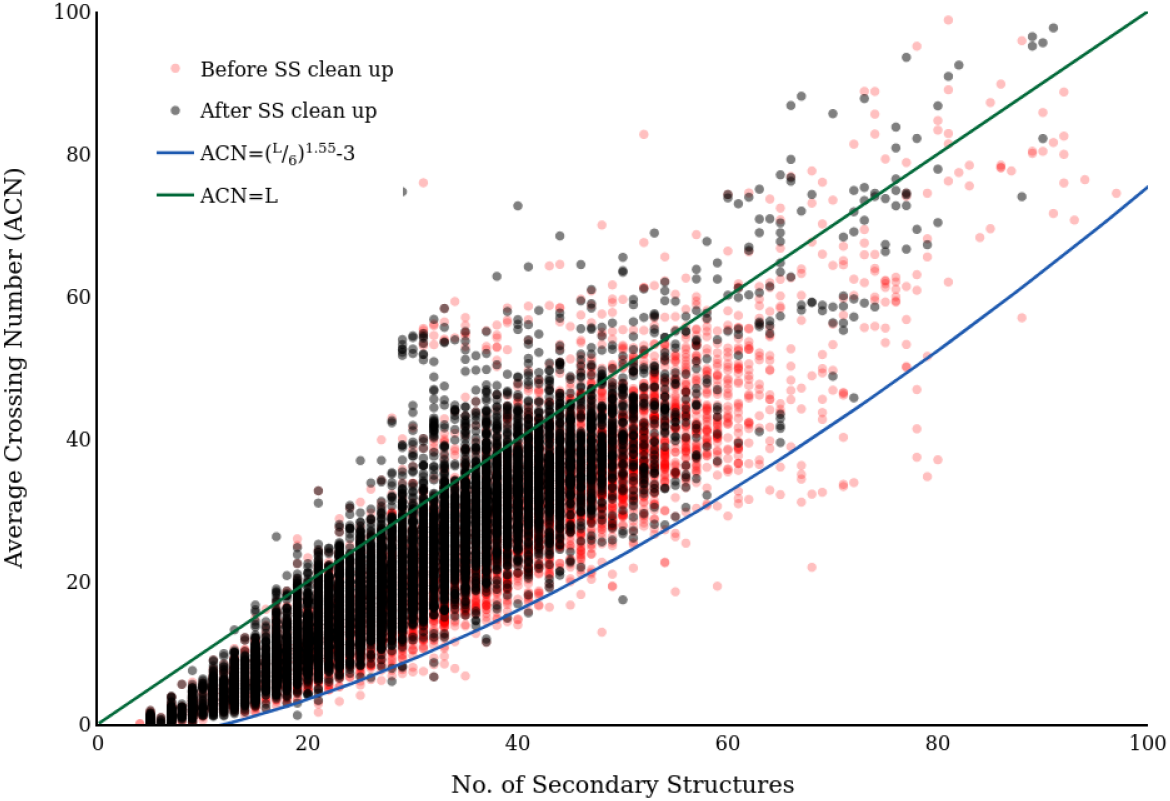
In red, the distribution of *acn* for the SKMT smoothed backbones of a representative sample of 10736 proteins from the PDB. In black, the distribution of *acn* for the same sample, where a clean up routine has been applied to the secondary structure prediction before computing the SKMT smoothed backbones.

There are a number of critical aspects of this plot we highlight.

1. 95.8% are bound from above by a linear growth of complexity *acn*(*C*) = *L*. This is commensurate with the fact we saw in the previous section a lot of complexity in protein structures arises from helical geometries (including the roadie type geometry).
2. 99.2% have an *acn* measure above the curve (*L/*6)^1.55^ –3, a fit obtained by sight. Implying a possible lower limit on the amount of complexity.
3. A curve (3*/*16)*L* log(*L*) roughly bisects the *acn* values above and below.

On the last curve, it was obtained in [84], where it is shown that *acn*(*C*) for an equilateral random walk of length *n* grows like *nlog*(*n*). This bound studied in the context of proteins in [62] where the *acn* of a sample of proteins is shown to follow this same length scaling. However, given our specific definition of length and that the smoothed protein backbone is clearly not equilateral, we can expect quite a different distribution of *acn* in our context. Nonetheless it is interesting it is roughly “in the middle” of the distribution. The fact there are many proteins which see a much faster, even super linear, growth in their *acn* with respect to their number of secondary structures implies some more systematic build up in complexity. Consider for example the signaling protein seen in Fig 18A, whose *acn* well exceeds not just *O*(*L* log *L*) but linear growth. It is clear that a random walk would be highly unlikely to achieve this specific *β*-barrel structure. On the other hand, the human filamin in Fig 18B has *acn* lying almost exactly on the *O*(*L* log *L*) curve. This immunoglobulin-like structure has many anti-parallel sheets, looking much more like the structure one might expect from a random walk.

### Falling below the empirical complexity

In order to use the *acn* in a predictive capacity, it is important to understand how strict this empirical lower bound is. In Fig 18C and D we highlight two of the proteins whose *acn* falls below the empirical bound displayed in blue. These represent the two most common reasons for a protein falling below the empirical bound on *acn*. Firstly, in Fig 9C we see a monomer unit of the trimeric Adenovirus Fibre. This protein forms a triple helix like structure as a trimer, which would have a large value of *acn*, however each single unit has relatively trivial entanglement. Since in this study we are only interested in monomers, for any multimer proteins that were pulled from the PDB we extracted just a single chain for analysis. This has little effect in terms of guiding our structural predictions, as we are mostly interested in resolving monomers. In the case we are aiming to predict a multimer, we can reduce the proportion of the penalty relative this bound. Fig 9D on the other hand shows a protein which has a structure that we would recognise as largely helical and systematic, it may be unclear then as to why this one falls below the empirical bound. This is due to the way in which we have defined length for this study. As the length is dependent on the number of secondary structural elements, there can be some anomalies when the secondary structure is poorly determined. In this case, the secondary structure prediction has split up the long linker sections between each of the yellow *β*-strands with some 1 amino acid long “*β*-strands”. As a result, our definition of length coinciding with the number of secondary structures sees this protein as much “longer” than it really is. Applying a simple routine removing any single amino acid long SSEs from the prediction, this protein falls well inside the blue bounding curve. These two examples are illustrative of all of the proteins which fall below this blue bounding curve. Indeed, in 19 we see that there remains only 0.14% of proteins falling below the blue curve. All of these remaining proteins are the relatively trivially folded single unit of a multimer structure, as in 18C.

This secondary structure “cleaning” is performed as standard in the SKMT algorithm, `simple_ss_clean`

### Using the ***acn*** to improve BioSAXS predictions

To see the effect of this *acn* bound on structural predictions, we consider the gene regulatory protein SMARCAL1. This protein regulates gene transcription through the alteration of the chromatin structure around those genes [89]. There is a predicted structure from AlphaFold for this protein, seen in Fig 20(a), however it has regions of low confidence, and most importantly is a poor fit to the SAXS data. This is illustrated in Figure Fig 20(b) where the SAXS scattering model is obtained using the method described in [24] (similar quality fits were obtained using the FOXS web server [90] as a check). For those readers not familiar with the BioSAXS data analysis, the factor *q* on the x-axis (inverse angstroms) measures the momentum transfer (the sine of the scattering angle divided by the x-ray wavelength) the vertical axis shows the logarithm of the observed scattering intensity *I*(*q*). The lower *q* range corresponds to larger scale structural information (overall shape *i.e.* elongated or globular) and the resolution increases with higher *q*. Fitting the low *q* range is of paramount importance as a small discrepancy with the data there can mean the overall shape of the molecule is wrong. By contrast at higher *q* (finer resolution) the experiment represents an average over a large number of molecules over a time period of approx 1s-1 min, so flexibility at that scale can mean discrepancies in a fixed prediction are less meaningful. Thus, for this simple illustrative example we stick to the *q* ∈ [0, 0.15] range. A rough rule of thumb for these experiments is that globular shapes have a hill shape low *q* SAXS curve whilst elongated structures a flatter low *q* (see *e.g.* Fig 3 of [91]) so Fig 20 indicates the AlphaFold prediction is too globular and the structure likely opens out somewhat in solution.

**Fig 20.**
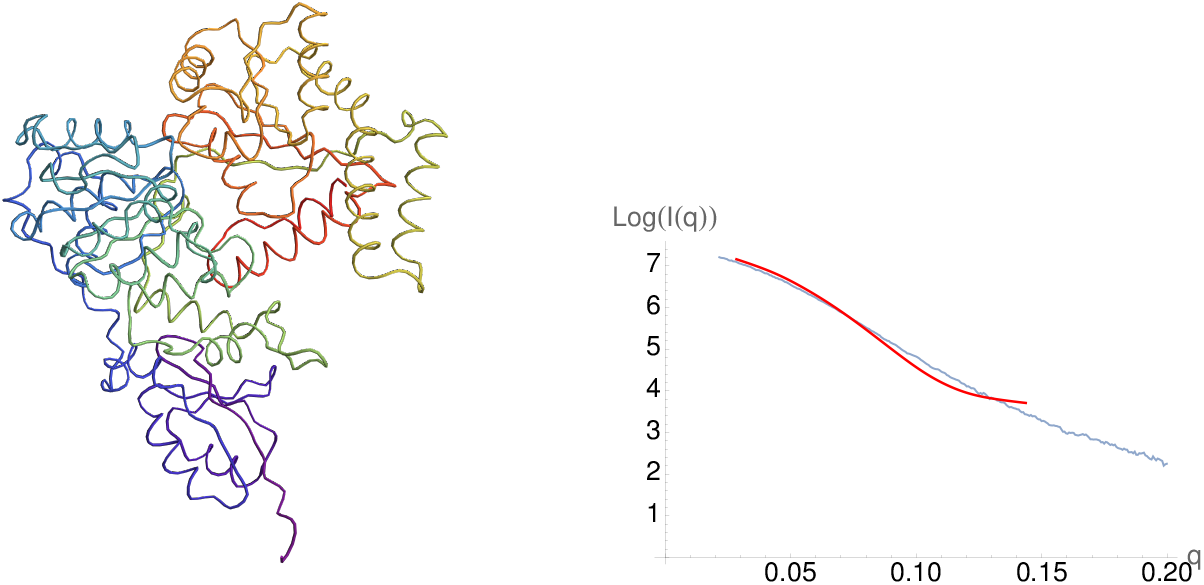
Panel (a) is the AlphaFold predicted structure for Human SMARCAL1. Panel (b) shows its fit (in red) to the scattering data (in blue) for *q ∈* [0, 0.15]

Using the constrained backbone algorithm [24], a new potential structure was produced to fit to the SAXS data. Briefly the method systematically varies the geometry of individual secondary structures of the C*^α^* backbone until the new structure fits the scattering data. It has local constraints on the C*^α^* backbone which were shown in [24] to mimic the Ramachandran constraints (so produces a locally plausible secondary structure) and a fitting penalty in order to prevent self-overlap of the molecule. What it does not have is any constraint on the viability, or the plausibility of the overall tertiary structure. This is a potential issue as the inverse scattering problem is not well posed [92] and multiple differing predictions for the data can be made. The viability of a prediction could in theory be tested using a full all atomistic physics model (and molecular dynamics simulations) but this is an extremely time consuming and intensive process. The fitting method introduced in [24] often needs to sample thousands of configurations in search so this would not be a practical possibility. By contrast the *acn* calculations are significantly quicker (of lower complexity than the scattering calculation itself) and the fact we have an apparent empirical lower bound means it can at the very least be used to rule out good fits to the scattering data which are unrealistically unfolded. We ran a series of fits to the SAXS data on *q* ∈ [0, 0.15] and calculated the *acn* of the final prediction. It should be clear this is not sufficient to make a clear prediction that the outcome is a plausible structure for the protein, further tests such as MD simulations and other experimental and statistical analysis would be needed for this. This is merely an example test of the potential efficacy of the *acn* measure as a means of providing a computationally efficient additional constraint on the tertiary fold search space.

Two examples are shown in Fig 21(a) and (c) to “open out” quite significantly. Visually we can see the three key subdomains of the structure are too far apart. Since SMARCAL1 has 84 secondary structure elements, we would expect its *acn* to be at least 56.8 (3.s.f) and likely a bit above this. The original structure’s value is 69.5 (3.s.f). This opened out structure however have an *acn* values 48.3 and 49.6 (both to 3.s.f). We then modified the constrained-backbone algorithm of [24] with a penalty for structures whose *acn* falls below the blue bounding curve (a simple step function for this testing exercises), we produce the prediction seen in Fig 22(a). This structure is much more globular although less so than the original structure). Its *acn* is 59.2 (3.s.f) which is above the bound.

**Fig 21.**
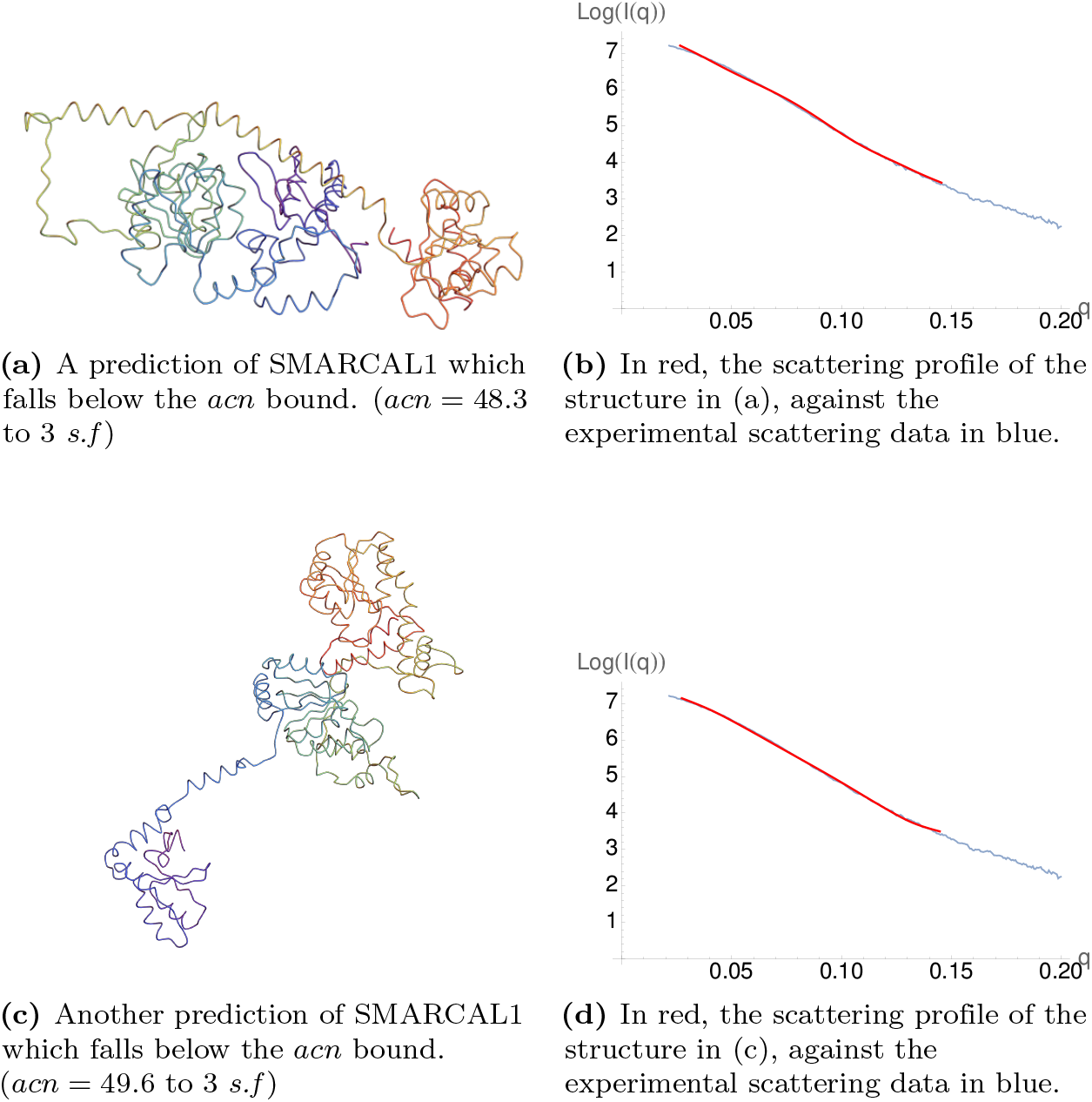
Examples of potential backbone structures which fit the SMARCAL1 data very well but are unrealistically unfolded according to the empirical bound on *acn*.

**Fig 22.**
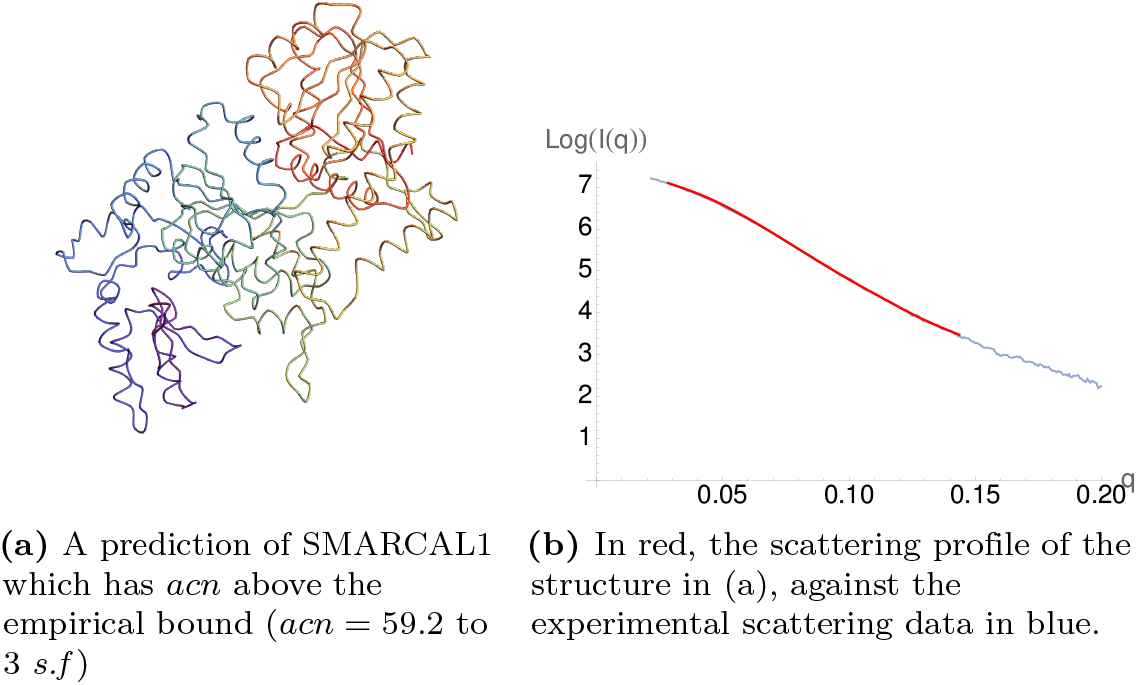
A predicted structure for SMARCAL1 which is both a good fit to the scattering data and is realistically folded according to the *acn*

As discussed earlier, this example is not supposed to be making any strong claims about the structure itself, here it is used as a convenient example. We argue this shows how this relatively simple to compute bound can be used in fold search space mechanisms as a practical means of enforcing some level of realism of the fold without the cost of including all the complex physics which go into a molecular dynamics model.

### Super-helical domains

Often when studying tertiary structure of proteins, we are interested in the folding of specific domains. For example, the CATH database provides information on protein domains that have a clear evolutionary link. These links are based on very specific folding motifs, with strict criteria on the number, length, and orientation of the secondary structural elements of the fold. One such example is the Rossmann fold [94], which consists of six *β*-strands, forming an extended *β* sheet, where the first three strands are connected via an *α* helix, giving an alternating *β – α – β – α –β* pattern. A schematic of this folding pattern is seen in Figure 23. It has been noted [95] that this initial alternating *α – β* segment is the most conserved aspect of the Rossmann fold. It is worth mentioning here that an alternating *α – β* motif is also seen in globally helical proteins such as that seen in Fig 12B.

**Fig 23.**
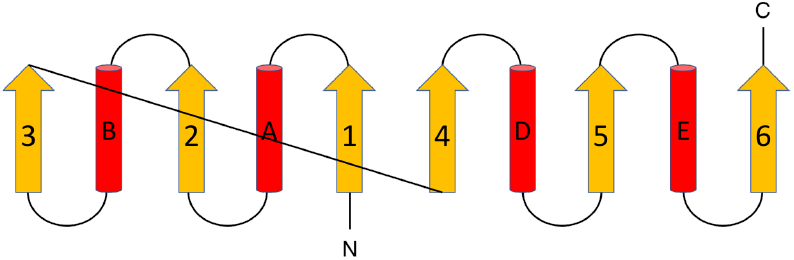
A schematic diagram of the six stranded Rossmann Fold. [93]

Crucially we have seen in our introductory example (Figure 2(d)) that this secondary motif leads to a helical super-geometry, which is similar to the motif for a TIM-barrel. A TIM-barrel domain consists of 8 *α*-helices and 8 parallel *β*-strands, which alternate along the backbone. The arrangement of the *β*-strands on the inside of the barrel is a key aspect of the stability for this structure [96]. Plots of *W* (_1*n*_) as a function of both curves are shown in Figure 24. One can see for the initial structure the overall gradient of growth is the same for both structures. We see between sections 5 and 42 the writhe in both cases grows to about 4.2 a gradient 0.12 (to 3*.s.f*)not too far off the gradient of 1.5 used for the linear curve in Figure 12.

**Fig 24.**
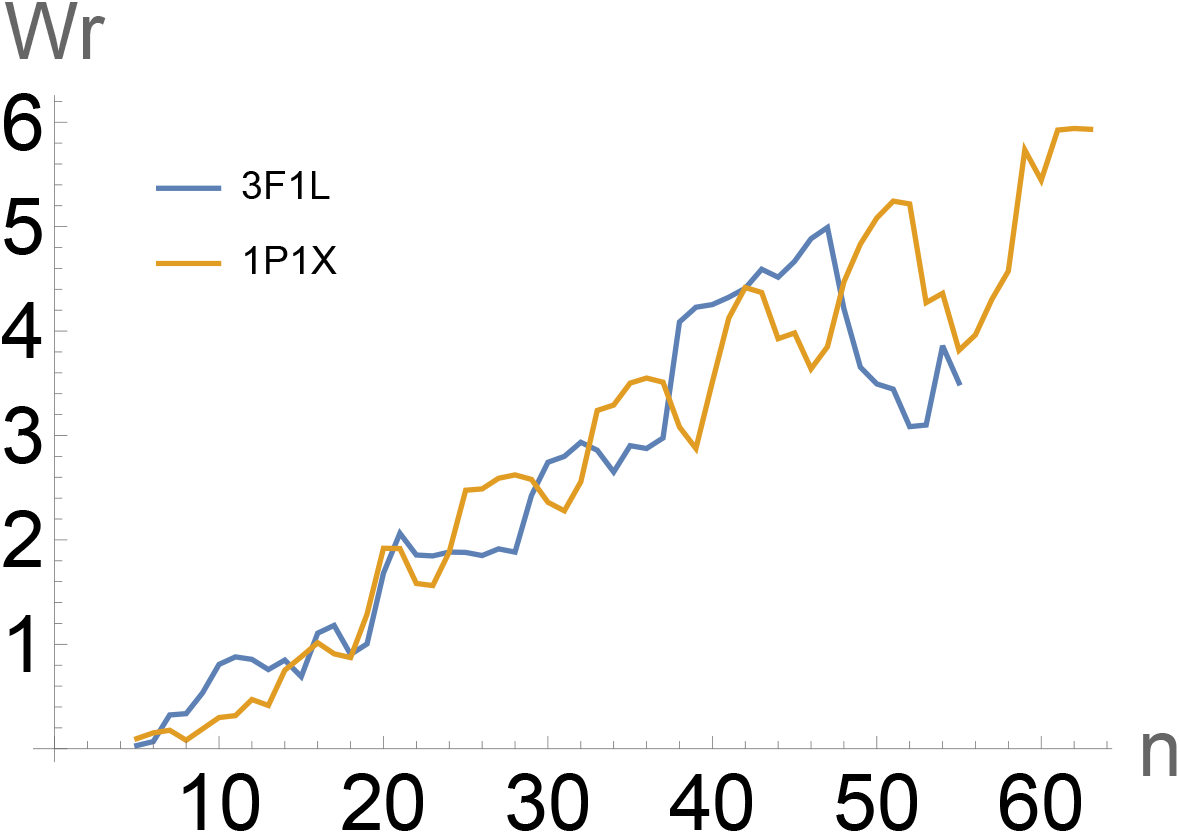
A comparison of the writhe profiles of two similarly helical protein structures. In orange, the TIM-Barrel domain 1P1X, in blue, the Rossmann fold domain 3F1L.

**Fig 25.**
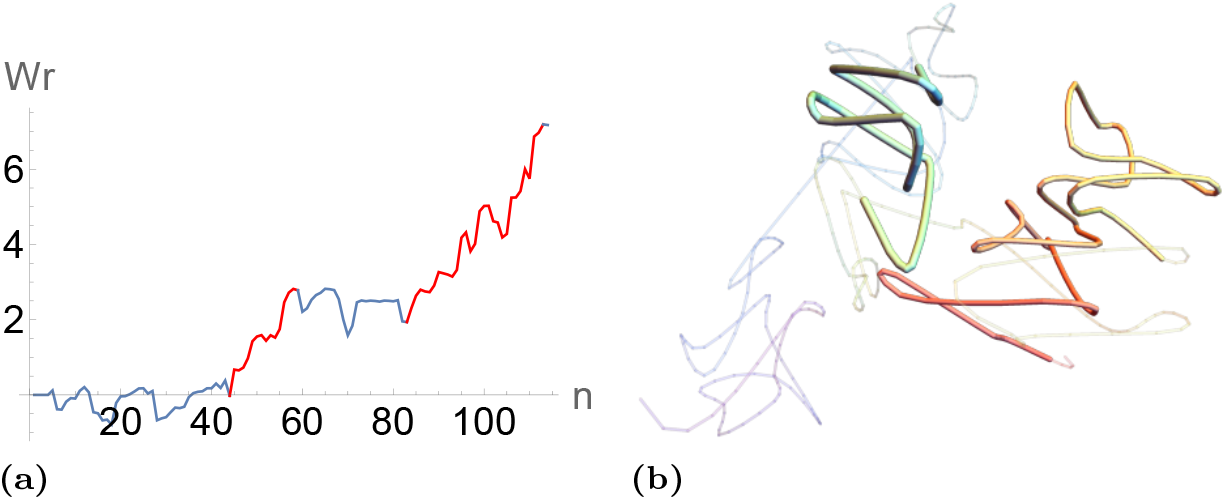
Panel (a), the writhe profile of the gene regulatory protein SMARCAL1. The subsections seeing a large linear growth in writhe are highlighted in red. Panel (b) the SKMT backbone curve. The two corresponding helical sections are shown opaquely against the rest of the structure to highlight their helical geometry.

Other proteins have helical subsections. We use the following subsection routine to identify helical. We consider subsets of the writhe profile of the SKMT smoothed structure subdomains *Wr*(*C_ij_*) of a reasonable length (*>*7 points). If the gradient of this subplot is greater than 0.1, we add it to a list of helical subsections. We then return the largest of all non overlapping such subsections. In the accompanying Colab notebook, one can include this method as an option when computing and plotting the writhe profile of your protein. An example of the output of this method is seen in 25 for the gene regulatory protein SMARCAL1 discussed in the previous section. As can be seen in this graph, there are two helical subdomains marked in red on the plot and on the (smoothed) structure. These two subdomains have a growth in writhe roughly equal to 0.2 times their length. The CATH “Search by Sequence” function predicts some sequence similarity between these subdomains to Rossmann fold domains. In order to study the subdomains more widely, we developed a routine which uses linear regression to reduce writhe plots to a few linear chunks, whose gradients can then be studied. This routine is described in more detail in the appendix. This routine was applied to our protein data set (the one described above), and the distribution these gradients is shown in Figure 26. We see a bimodal distribution the bulk of whose values lie between 0.06 and 0.2 in absolute value. This helical search can be applied to a protein using the `highlight_helical_subsections` option when plotting the writhe in the SWRITHE notebook.

**Fig 26.**
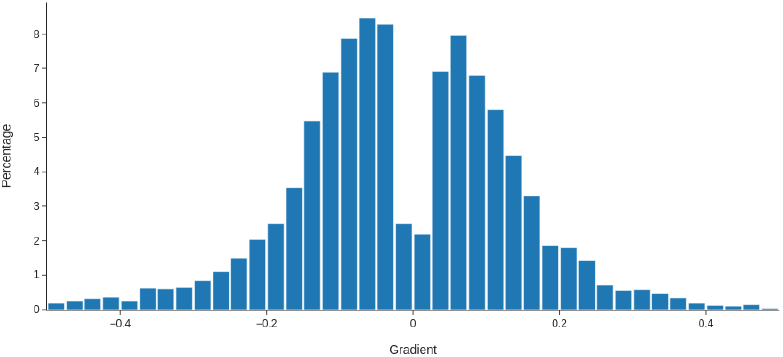
The distribution of gradients for linear subsections of the writhe profiles of SKMT smooth backbones from our representative sample of the PDB.

### Comparing structures using *W* (*C_ij_*)

The apparent similarity between the TIM barrel and Rossmann fold examples of the previous section opens the question as to whether we can identify any other surprising tertiary structural similarities by comparing writhe development distributions. To test this we create the following similarity metric for subsections 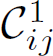 and 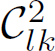 of (SKMT smoothed) curves *C*^1^*, C*^2^, where *l − k* = *j − i*, so that the regions compared are the same size (but we allow for example that one structure could be a subdomain of another). We measure their similarity *S*(*C*^1^ *, C*^2^)as follows:

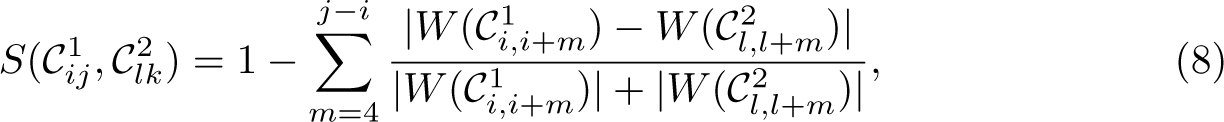

which measures the difference in writhe of all subsections [*i, i* + *m*], [*l, l* + *m*] weighted by the writhe values themselves; its value is between [0, 1]. This metric is applied to all similar size subsections of the two structures (above say *L* = 8) and the Largest length sections score for which *S >* 0.85 is recorded 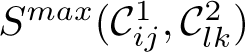. For reference *S^max^* for the structures 1P1X (Tim barrel) and 3F1L (Rossmann fold) is 0.85 (2.s.f) for a section of length 50: practically all of the structure 1P1X. There are subsections with higher comparison values of *S* but these had smaller lengths. The metric 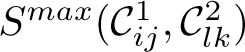 can be calculated for two structures using the `find_sim_sections` function in the SWRITHE library.

We applied this comparison metric 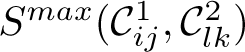 for the structure 3F1L (a Rossmann fold) and all others in our database (the 10736 used in the previous sections). A large number had a length 0 (no structural similarities at all) but there are often a good number of small length similarities. For now we restrict to structures for which 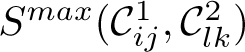 *>* 0.85 for *L >* 50 so we are effectively comparing similarity to the whole backbone of 3F1L. We find 151 such cases. Their writhe plots are shown in Fig 27. A comparison to the CATH classification of these proteins show 81.1% of them were classed as Rossmann folds (as one would expect) 8.3% were TIM barrel proteins seemingly strengthening the apparent structural similarity relationship between these fold types. The matched data set is available in the supplementary material (`Simto3f1l.txt`).

**Fig 27.**
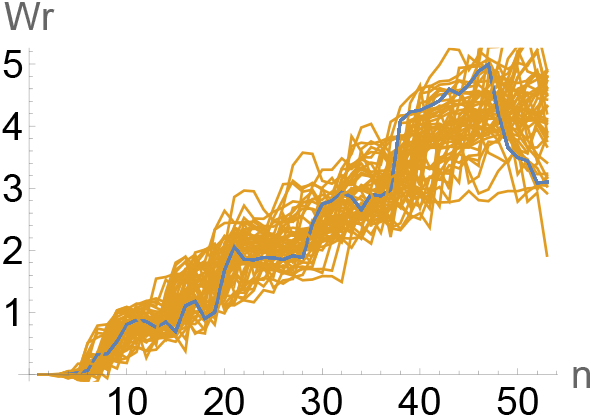
In blue, a plot of the writhe of the SKMT smoothed backbone of 3F1L. In orange, the writhe profiles of all SKMT smoothed backbones that are classed as similar by the *S^max^* metric. 8

We performed a similar comparison for 1P1X to our database. This yielded 232 structures of which 63.3% were Rossmann folds, and 25.5% TIM barrels. For context the full database has a share of 18.3% Rossmann folds and 3.4% TIM barrels. So there seems to be a very strong similarity of tertiary geometry between these two families of structures. There is a small number of other CATH classified proteins which also matched this structure. We do not analyse the similarity here leaving it for future study. The matched data set is available in the supplementary material (`Simto1p1x.txt`).

### Roadie like geometries

Applying our roadie criteria for subsections: *Wr*(*C_in_*) *<* 0.05 and for some *Wr*(*_im_*)*, m < n* we have *Wr*(*_im_*) *>* 0.90, to our database we found 2950 examples, nearly 30%. The set of proteins with markedly roadie like geometry in all or part of their structure is available in the supplementary material (`RoadieList.txt`). Some examples are shown in Fig 28. The Viral Chemokine is particuarkly striking as the roadie geometry essentially accounts for the entire structure. This check can be applied to a structure using the `highlight_roadie_sections` option when plotting the writhe profile of a protein. Compared to the helical comparisons above, these Roadie like proteins show no real trend in relation to the CATH classification, as can be seen in Fig 29. Though this set may again seem to be dominated by Rossmann folds, this is more due to their large representation in our full data set than a true link to Roadie geometries. Besides this, there is no clear tendency to a specific CATH topology for these structures. The Roadie-like structure is identified via strict writhe criteria, meaning this a truly systematic method of entanglement for subdomains. The physical uses for this structure are clear in other contexts, whether this is applicable to proteins though is beyond the scope of this study but we present this method and data so that the structural biology community might identify a reason for its prevalence.

**Fig 28.**
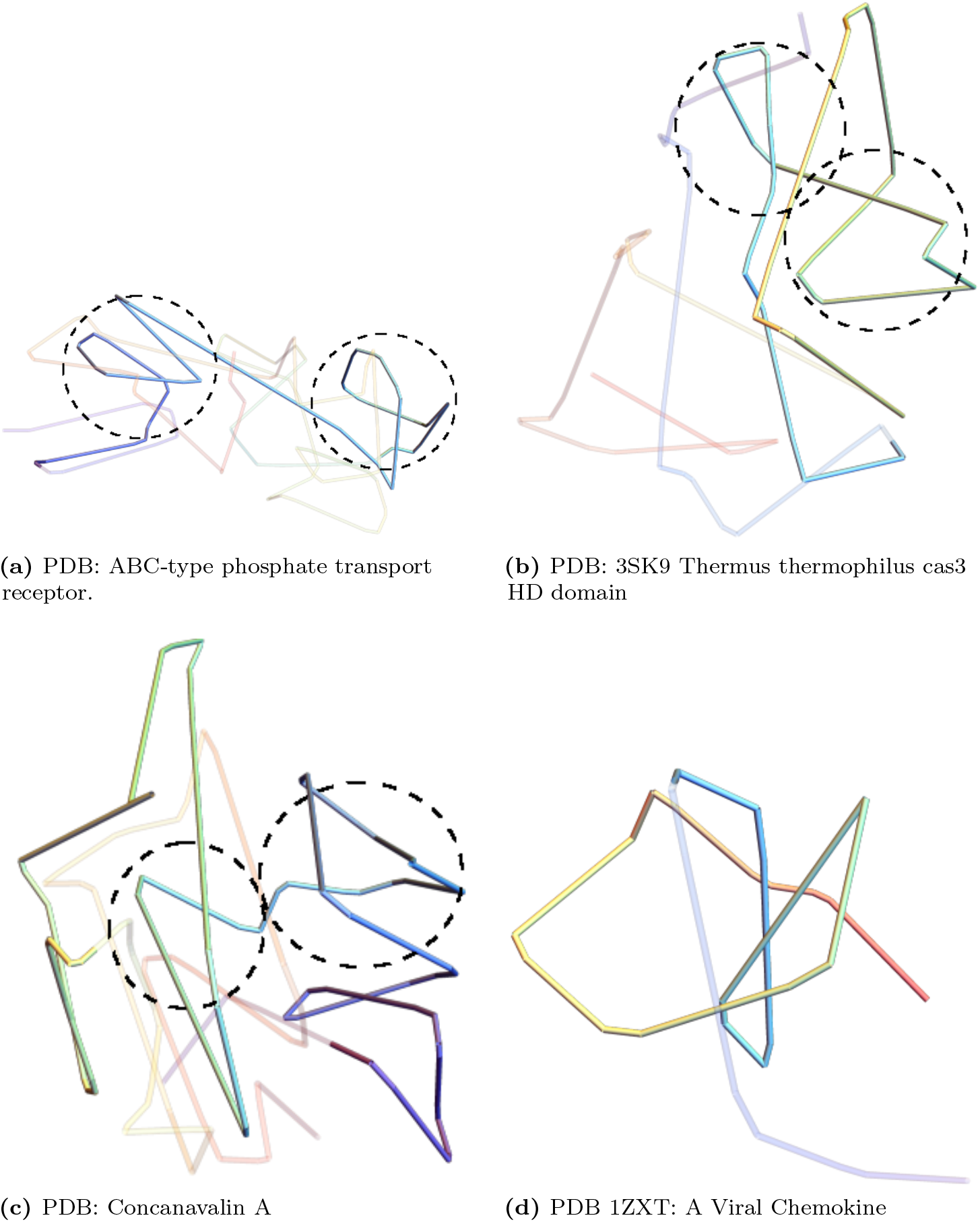
Examples of roadie (net zero writhe) geometry found in our PDB data set.

**Fig 29.**
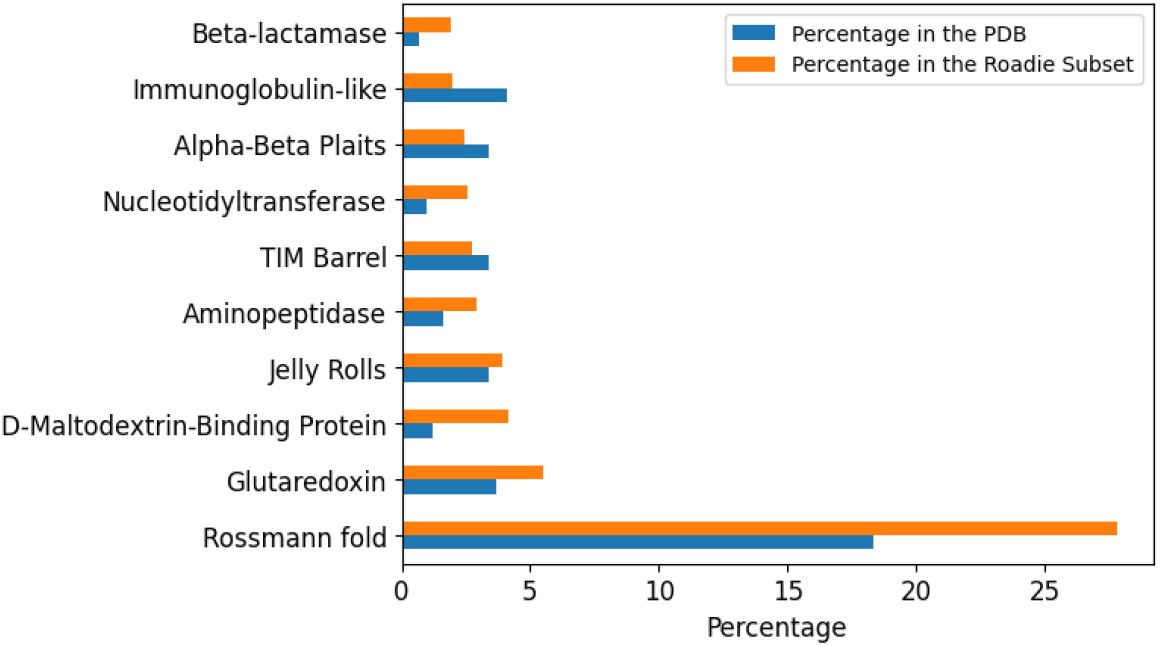
A histogram of the 10 highest represented CATH topologies in the set of Roadie like proteins in orange. The percentage of these topologies in the general data set is shown in blue for comparison. IN the first three cases the two counter loops are encircled. In case (d) the roadie geometry is essentially the entire protein.

## Discussion/Conclusion

In this study, we detail a series of metrics and methods for utilising the writhe *Wr* and average crossing number *acn* as a measure of tertiary structure entanglement for protein backbone curves. First, we highlight how fundamental properties of the writhe such as linear growth of helical geometries, allow us to find similarities in proteins which other methods may miss. Second, we apply it to a smoothed backbone structure, which treats secondary structures such as alpha helices and strand helices as rigid units and simplify linker sections whilst preserving their fundamental topology of the structure. We name this smoothing (coarse-graining) protocol the SKMT algorithm: secondary KMT, were the KMT algorithm is a known method to preserve topology whilst simplifying a structure. This we show how it can be used to asses the topological changes (changes in connectivity) of similar or evolving structures.

In this study, we have shown that the writhe of protein backbone curves is bounded with respect to the number of secondary structure elements it has. This is done by applying the metric to a set of *>* 10000 sequentially unique (at the 60% level) high quality structures. In particular, a linear bounding curve of growth in the writhe with respect to the number of secondary structures, with a gradient of 0.15, contains 97.4% of protein structures. We show that this linear scaling is directly linked to the geometry of globally helical protein structures. This same helical geometry is consistently found in sub-sections of other proteins. This helical geometry (on the tertiary not secondary scale) is a method of systematically building up writhe in such a way as to produce thermally stable structures with strong inter sheet bonds (thermally stable TIM-barrel structures being a common example). By comparing this to the distribution of writhe for trefoil knotted proteins, we see that there is no real correlation between knottedness and writhe. Conversely, many proteins exhibit zero net entanglement whilst exhibiting significant complexity, across all length scales. Here we identify “roadie” like protein substructures which have no net writhe but are significantly coiled. These substructures, in systematic way, build up entanglement and then undo it, mimicking the process by which cables can be folded without knotting themselves. Though the functional explanation for this structure is not yet clear, it is commonly seen across many families of proteins and has many uses as a conformation in other disciplines.

We then study the distribution of unsigned writhing, or total complexity *acn* for proteins, again establishing empirical bounds on this measure of complexity as a function of the number of secondary sections of the protein. We show again that linear growth is a good upper bound on complexity, strengthening the case that complexity of entanglement in proteins is due to large scale helical geometries. We also provide an empirical lower bound on the *acn* proteins, which we show is effective when used as a constraint for realistic and efficient searches of potential structure landscapes. An example of how this can be applied to BioSAXS data is detailed.

By studying the writhe profile of subdomains of proteins we identify these large scale helical geometries exist not just as global conformations, but are also present as part of larger complexes. These structures are especially present in Rossmann folds and TIM barrels, which highlights the link to thermal stability. We show there is often a significant similarity of fold geometry between these structure classifications which is not readily revealed by other structural comparison methods. The writhe measure is also able to identify roadie-like substructures in proteins, these are significantly entangled structures which exhibit coiling of opposing chirality (again on the tertiary not secondary scale). Though the functional advantage of these specific geometries is not yet clear, they are ubiquitous across many families of proteins.

Alongside this work we provide a Colab notebook where one can

1. Apply the SKMT secondary backbone smoothing algorithm to a specified PDB structure, using the skmt function.
2. Compute the writhe *Wr* and average crossing number *acn* of the smoothed backbone of a given protein using the `calculate_writhe` and `calculate_abs_writhe` functions.
3. Highlight helical subsections of a given structure by selecting the highlight helical subsections option when plotting the writhe profile.
4. Identify subsections of two proteins of similar geometry through the metric *S^max^* a measure of the largest subsection of the two proteins which have significantly similar writhe distributions, using the `find_sim_sections` function.
5. Identify roadie geometries, subsections with significant complexity (*acn*) but no net writhe, by selecting the `highlight_roadie_sections` option when plotting the writhe profile.

## Supporting information

### S1 - Identifying linear subsections of writhe plots

The routine used to subdivide writhe plots into linear subsections is as follows.

1. We first perform a LOWESS (locally weighted scatterplot smoothing) on the writhe data.
2. Perform a linear regression on the writhe graph. If the coefficient of determination (*R*^2^) is greater than 0.75, we accept this fit.
3. Otherwise, for *k* = 4, &, *n* 4 where *n* is the max *x* value of the writhe graph, we split the graph into two parts either side of *k*. We again perform a linear regression on these two subplots, average their *R*^2^ values, and store them. If the max of the averaged *R*^2^ values is greater than 0.75, we accept these two linear subdomains.
4. Otherwise, we pick two points *k* = 4, &, *n* 8, *l* = 8, &, *n* 4 and consider the three subplots defined by these splitting points. As above, we perform linear regression on the respective subplots, and store the averaged *R*^2^. We accept if the max is greater then 0.75.
5. Otherwise, consider three splitting points to give four subplots. Scoring and accepting as before.
6. Finally, if none of the above fits are above our tolerance, we consider all subplots with at least 4 points, and find the best fitting linear regression of these, accepting if its *R*^2^ is greater than 0.75.

## Acknowledgments

This work was funded by the EPSRC MoSMed CDT (EP/S022791/1, studentship award to AB) and Diamond Light Source (studentship award to AB).

## References

1. Diamond R. Real-space refinement of the structure of hen egg-white lysozyme. Journal of Molecular Biology. 1974;82(3):371–391. doi:https://doi.org/10.1016/0022-2836(74)90598-1.

2. Otaki JM, Tsutsumi M, Gotoh T, Yamamoto H. Secondary structure characterization based on amino acid composition and availability in proteins. Journal of chemical information and modeling. 2010;50(4):690–700.

3. Orengo CA, Todd AE, Thornton JM. From protein structure to function. Current Opinion in Structural Biology. 1999;9(3):374–382. doi:https://doi.org/10.1016/S0959-440X(99)80051-7.

4. Berman HM, Westbrook J, Feng Z, Gilliland G, Bhat TN, Weissig H, et al. The Protein Data Bank. Nucleic Acids Research. 2000;28(1):235–242. doi:10.1093/nar/28.1.235.

5. Jumper J, Evans R, Pritzel A, Green T, Figurnov M, Ronneberger O, et al. Highly accurate protein structure prediction with AlphaFold. Nature. 2021;596(7873):583–589.

6. Baek M, DiMaio F, Anishchenko I, Dauparas J, Ovchinnikov S, Lee GR, et al. Accurate prediction of protein structures and interactions using a three-track neural network. Science. 2021;373(6557):871–876.

7. Sillitoe I, Bordin N, Dawson N, Waman VP, Ashford P, Scholes HM, et al. CATH: increased structural coverage of functional space. Nucleic Acids Research. 2020;49(D1):D266–D273. doi:10.1093/nar/gkaa1079.

8. Andreeva A, Kulesha E, Gough J, Murzin AG. The SCOP database in 2020: expanded classification of representative family and superfamily domains of known protein structures. Nucleic Acids Research. 2019;48(D1):D376–D382. doi:10.1093/nar/gkz1064.

9. Holm L. Dali server: structural unification of protein families. Nucleic Acids Research. 2022;50(W1):W210–W215. doi:10.1093/nar/gkac387.

10. Orengo CA, Pearl FMG, Bray JE, Todd AE, Martin AC, Lo Conte L, et al. The CATH Database provides insights into protein structure/function relationships. Nucleic Acids Research. 1999;27(1):275–279. doi:10.1093/nar/27.1.275.

11. Heine A, Luz JG, Wong CH, Wilson IA. Analysis of the Class I Aldolase Binding Site Architecture Based on the Crystal Structure of 2-Deoxyribose-5-phosphate Aldolase at 0.99Å Resolution. Journal of Molecular Biology. 2004;343(4):1019–1034. doi:https://doi.org/10.1016/j.jmb.2004.08.066.

12. Vijayalakshmi J, Meredith TC, Woodard RW. The 0.95 A structure of an oxidoreductase, yciK from E.coli. None. 2008;.

13. Akdel M, Pires DE, Pardo EP, Jänes J, Zalevsky AO, Mésźaros B, et al. A structural biology community assessment of AlphaFold2 applications. Nature Structural & Molecular Biology. 2022; p. 1–12.

14. Figueroa Ýevenes M, Sleutel M, Vandevenne M, Parvizi G, Attout S, Jacquin O, et al. The unexpected structure of the designed protein Octarellin V.1 forms a challenge for protein structure prediction tools. Journal of Structural Biology. 2016;doi:10.1016/j.jsb.2016.05.004.

15. Rambo RP, Tainer JA. Super-resolution in solution X-ray scattering and its applications to structural systems biology. Annual review of biophysics. 2013;42:415–441.

16. Mertens HD, Svergun DI. Structural characterization of proteins and complexes using small-angle X-ray solution scattering. Journal of structural biology. 2010;172(1):128–141.

17. Kikhney AG, Svergun DI. A practical guide to small angle X-ray scattering (SAXS) of flexible and intrinsically disordered proteins. FEBS letters. 2015;589(19):2570–2577.

18. Schneidman-Duhovny D, Hammel M. Modeling structure and dynamics of protein complexes with SAXS profiles. Springer; 2018.

19. Castellví A, Pascual-Izarra C, Crosas E, Malfois M, Juanhuix J. Improving data quality and expanding BioSAXS experiments to low-molecular-weight and low-concentration protein samples. Acta Crystallographica Section D: Structural Biology. 2020;76(10):971–981.

20. Osz J, McEwen AG, Bourguet M, Przybilla F, Peluso-Iltis C, Poussin-Courmontagne P, et al. Structural basis for DNA recognition and allosteric control of the retinoic acid receptors RAR–RXR. Nucleic Acids Research. 2020;48(17):9969–9985. doi:10.1093/nar/gkaa697.

21. Vestergaard B, Sanyal S, Roessle M, Mora L, Buckingham RH, Kastrup JS, et al. The SAXS solution structure of RF1 differs from its crystal structure and is similar to its ribosome bound cryo-EM structure. Molecular cell. 2005;20(6):929–938.

22. Hura GL, Hodge CD, Rosenberg D, Guzenko D, Duarte JM, Monastyrskyy B, et al. Small angle X-ray scattering-assisted protein structure prediction in CASP13 and emergence of solution structure differences. Proteins: Structure, Function, and Bioinformatics. 2019;87(12):1298–1314.

23. Rambo RP, Tainer JA. Accurate assessment of mass, models and resolution by small-angle scattering. Nature. 2013;496(7446):477–481.

24. Prior C, Davies OR, Bruce D, Pohl E. Obtaining Tertiary Protein Structures by the ab Initio Interpretation of Small Angle X-ray Scattering Data. Journal of Chemical Theory and Computation. 2020;16(3):1985–2001. doi:10.1021/acs.jctc.9b01010.

25. Sadowski MI. In: Roberts GCK, editor. Protein Structure Comparison Methods. Berlin, Heidelberg: Springer Berlin Heidelberg; 2013. p. 2055–2060. Available from: https://doi.org/10.1007/978-3-642-16712-6_413.

26. Zhang Y, Skolnick J. Scoring function for automated assessment of protein structure template quality. Proteins: Structure. 2004;57.

27. Gelly JC, Joseph AP, Srinivasan N, de Brevern AG. iPBA: a tool for protein structure comparison using sequence alignment strategies. Nucleic Acids Research. 2011;39:W18–W23. doi:10.1093/nar/gkr333.

28. Zhang Y, Skolnick J. TM-align: a protein structure alignment algorithm based on the TM-score. Nucleic Acids Research. 2005;33(7):2302–2309. doi:10.1093/nar/gki524.

29. Li Z, Jaroszewski L, Iyer M, Sedova M, Godzik A. FATCAT 2.0: towards a better understanding of the structural diversity of proteins. Nucleic Acids Research. 2020;48(W1):W60–W64. doi:10.1093/nar/gkaa443.

30. Mansfield M. Are there knots in proteins? Nature structural biology. 1994;1:213–4. doi:10.1038/nsb0494-213.

31. Dabrowski-Tumanski P, Sulkowska JI. Topological knots and links in proteins. Proceedings of the National Academy of Sciences. 2017;114(13):3415–3420. doi:10.1073/pnas.1615862114.

32. Virnau P, Mirny LA, Kardar M. Intricate knots in proteins: Function and evolution. PLoS computational biology. 2006;2(9):e122.

33. Soler MA, Faisca PF. Effects of knots on protein folding properties. PloS one. 2013;8(9):e74755.

34. Dabrowski-Tumanski P, Stasiak A, Sulkowska JI. In search of functional advantages of knots in proteins. PloS one. 2016;11(11):e0165986.

35. Lua RC, Grosberg AY. Statistics of Knots, Geometry of Conformations, and Evolution of Proteins. PLOS Computational Biology. 2006;2(5):1–8. doi:10.1371/journal.pcbi.0020045.

36. King NP, Yeates EO, Yeates TO. Identification of Rare Slipknots in Proteins and Their Implications for Stability and Folding. Journal of Molecular Biology. 2007;373(1):153–166. doi:https://doi.org/10.1016/j.jmb.2007.07.042.

37. Turaev V. Knotoids. Osaka J Math. 2012;49:195–223.

38. Goundaroulis D, Dorier J, Benedetti F, Stasiak A. Studies of global and local entanglements of individual protein chains using the concept of knotoids. Scientific reports. 2017;7(1):6309.

39. Barbensi A, Yerolemou N, Vipond O, Mahler BI, Dabrowski-Tumanski P, Goundaroulis D. A Topological Selection of Folding Pathways from Native States of Knotted Proteins. Symmetry. 2021;13(9). doi:10.3390/sym13091670.

40. Koniaris K, Muthukumar M. Self-entanglement in ring polymers. The Journal of Chemical Physics. 1991;95(4):2873–2881. doi:10.1063/1.460889.

41. Klenin K, Langowski J. Computation of writhe in modeling of supercoiled DNA. Biopolymers. 2000;54(5):307–317. doi:https://doi.org/10.1002/1097-0282(20001015)54:5¡307::AID-BIP20¿3.0.CO;2-Y.

42. Dennis M, Hannay J. Geometry of Căluăreanu’s theorem. Proceedings of the Royal Society A: Mathematical, Physical and Engineering Sciences. 2005;461(2062):3245–3254.

43. Klenin K, Vologodskii A, Anshelevich V, Klishko VY, Dykhne A, Frank-Kamenetskii M. Variance of writhe for wormlike DNA rings with excluded volume. Journal of Biomolecular Structure and Dynamics. 1989;6(4):707–714.

44. Fain B, Rudnick J, Östlund S. Conformations of linear DNA. Physical Review E. 1997;55(6):7364.

45. Bouchiat C, Mezard M. Elasticity model of a supercoiled DNA molecule. Physical review letters. 1998;80(7):1556.

46. Marko JF, Neukirch S. Competition between curls and plectonemes near the buckling transition of stretched supercoiled DNA. Physical Review E. 2012;85(1):011908.

47. Lam PM, Zhen Y. Twisting, supercoiling and stretching in protein bound DNA. Physica A: Statistical Mechanics and its Applications. 2018;496:200–208.

48. Sierzega Z, Wereszczynski J, Prior C. WASP: a software package for correctly characterizing the topological development of ribbon structures. Scientific Reports. 2021;11(1):1527.

49. Marenduzzo D, Micheletti C, Orlandini E, Sumners DW. Topological friction strongly affects viral DNA ejection. Proceedings of the National Academy of Sciences. 2013;110(50):20081–20086. doi:10.1073/pnas.1306601110.

50. Bednar J, Furrer P, Stasiak A, Dubochet J, Egelman EH, Bates AD. The twist, writhe and overall shape of supercoiled DNA change during counterion-induced transition from a loosely to a tightly interwound superhelix: possible implications for DNA structure in vivo. Journal of molecular biology. 1994;235(3):825–847.

51. Arsuaga J, Vazquez M, McGuirk P, Trigueros S, Sumners DW, Roca J. DNA knots reveal a chiral organization of DNA in phage capsids. Proceedings of the National Academy of Sciences. 2005;102(26):9165–9169.

52. Sumners D. The role of knot theory in DNA research. In: Geometry and Topology. CRC Press; 2020. p. 297–318.

53. Sleiman JL, Burton RH, Caraglio M, Gutierrez Fosado YA, Michieletto D. Geometric Predictors of Knotted and Linked Arcs. ACS Polymers Au. 2022;2(5):341–350.

54. Røgen P, Fain B. Automatic Classification of Protein Structure by Using Gauss Integrals. Proceedings of the National Academy of Sciences of the United States of America. 2003;100(1):119–124.

55. Røgen P, Bohr H. A new family of global protein shape descriptors. Mathematical Biosciences. 2003;182(2):167–181. doi:https://doi.org/10.1016/S0025-5564(02)00216-X.

56. Chang PL, Rinne AW, Dewey TG. Structure alignment based on coding of local geometric measures. BMC bioinformatics. 2006;7(1):1–10.

57. Zhi D, Shatsky M, Brenner SE. Alignment-free local structural search by writhe decomposition. Bioinformatics. 2010;26(9):1176–1184.

58. Grønbæk C, Hamelryck T, Røgen P. GISA: using Gauss Integrals to identify rare conformations in protein structures. PeerJ. 2020;8:e9159. doi:10.7717/peerj.9159.

59. Cantarella J, DeTurck D, Gluck H. Upper bounds for the writhing of knots and the helicity of vector fields. AMS IP Studies in Advanced Mathematics. 2001;24:1–22.

60. Rose GD, Fleming PJ, Banavar JR, Maritan A. A backbone-based theory of protein folding. Proceedings of the National Academy of Sciences. 2006;103(45):16623–16633.

61. Panagiotou E, Millett KC, Lambropoulou S. The linking number and the writhe of uniform random walks and polygons in confined spaces. Journal of Physics A: Mathematical and Theoretical. 2010;43(4):045208. doi:10.1088/1751-8113/43/4/045208.

62. Dobay A, Dubochet J, Stasiak A, Diao Y. In: Scaling of the Average Crossing Number in Equilateral Random Walks, Knots and Proteins. World Scientific; 2005. p. 219–231. Available from: https://www.worldscientific.com/doi/abs/10.1142/9789812703460_0012.

63. Ramakrishnan C, Balasubramanian R. Stereochemical Criteria for Polypeptide and Protein Chain Conformations. International Journal of Peptide and Protein Research. 1972;4(2):79–90. doi:https://doi.org/10.1111/j.1399-3011.1972.tb03403.x.

64. Arteca GA. Scaling regimes of molecular size and self-entanglements in very compact proteins. Physical Review E. 1995;51(3):2600.

65. Klenin K, Langowski J. Computation of writhe in modeling of supercoiled DNA. Biopolymers. 2000;54(5):307–317.

66. Wollert T, Heinz DW, Schubert WD. Thermodynamically reengineering the listerial invasion complex InlA/E-cadherin. Proceedings of the National Academy of Sciences. 2007;104(35):13960–13965.

67. White JH, Bauer WR. Calculation of the twist and the writhe for representative models of DNA. Journal of molecular biology. 1986;189(2):329–341.

68. Majorek KA, Porebski PJ, Dayal A, Zimmerman MD, Jablonska K, Stewart AJ, et al. Structural and immunologic characterization of bovine, horse, and rabbit serum albumins. Molecular immunology. 2012;52(3-4):174–182.

69. org AH. Repairs: Coiling Cables; 2022. Available from: http://audio.hortonwho.org/reinforcement/repairs/repairs.htm.

70. Porta J, Kolar C, Kozmin SG, Pavlov YI, Borgstahl GE. Structure of the orthorhombic form of human inosine triphosphate pyrophosphatase. Acta Crystallographica Section F: Structural Biology and Crystallization Communications. 2006;62(11):1076–1081.

71. Singh R, Hawkins W. Sutures, ligatures and knots. Surgery (Oxford). 2017;35(4):185–189.

72. Prior CB, Neukirch S. The extended polar writhe: a tool for open curves mechanics. Journal of Physics A: Mathematical and Theoretical. 2016;49(21):215201.

73. Benjamin K, Mukta L, Moryoussef G, Uren C, Harrington HA, Tillmann U, et al.. Homology of homologous knotted proteins; 2022. Available from: https://arxiv.org/abs/2201.07709.

74. Dabrowski-Tumanski P, Stasiak A, Sulkowska J. In Search of Functional Advantages of Knots in Proteins. PLOS ONE. 2016;11. doi:10.1371/journal.pone.0165986.

75. Sulkowska JI. On folding of entangled proteins: knots, lassos, links and *θ*-curves. Current opinion in structural biology. 2020;60:131–141.

76. Shi D, Morizono H, Yu X, Roth L, Caldovic L, Allewell NM, et al. Crystal structure of N-acetylornithine transcarbamylase from Xanthomonas campestris: a novel enzyme in a new arginine biosynthetic pathway found in several eubacteria. Journal of Biological Chemistry. 2005;280(15):14366–14369.

77. Barta ML, Thomas K, Yuan H, Lovell S, Battaile KP, Schramm VL, et al. Structural and Biochemical Characterization of Chlamydia trachomatis Hypothetical Protein CT263 Supports That Menaquinone Synthesis Occurs through the Futalosine Pathway*. Journal of Biological Chemistry. 2014;289(46):32214–32229. doi:https://doi.org/10.1074/jbc.M114.594325.

78. Wollert T, Heinz DW, Schubert WD. Thermodynamically reengineering the listerial invasion complex InlA/E-cadherin. Proceedings of the National Academy of Sciences. 2007;104(35):13960–13965. doi:10.1073/pnas.0702199104.

79. for Structural Genomics (JCSG) JC. Crystal structure of Putative DNA-binding protein (*Y P*_2_99413.1) from Ralstonia eutrophA JMP134 at 1.30 A resolution. None. 2009;.

80. Wang H, DeRose E, London R, Shears S. IP6K Structure and the Molecular Determinants of Catalytic Specificity in an Inositol Phosphate Kinase Family. Nature communications. 2014;5:4178. doi:10.1038/ncomms5178.

81. Dorier J, Goundaroulis D, Benedetti F, Stasiak A. Knoto-ID: a tool to study the entanglement of open protein chains using the concept of knotoids. Bioinformatics. 2018;34(19):3402–3404. doi:10.1093/bioinformatics/bty365.

82. Banavar JR, Maritan A, Micheletti C, Trovato A. Geometry and physics of proteins. Proteins: Structure, Function, and Bioinformatics. 2002;47(3):315–322.

83. Barbensi A, Yerolemou N, Vipond O, Mahler B, Dabrowski-Tumanski P, Goundaroulis D. A Topological Selection of Folding Pathways from Native States of Knotted Proteins. Symmetry. 2021;13:1670. doi:10.3390/sym13091670.

84. Diao Y, Dobay A, Kusner RB, Millett K, Stasiak A. The average crossing number of equilateral random polygons. Journal of Physics A: Mathematical and General. 2003;36(46):11561–11574. doi:10.1088/0305-4470/36/46/002.

85. Wang Q, Shui B, Kotlikoff MI, Sondermann H. Structural Basis for Calcium Sensing by GCaMP2. Structure. 2008;16(12):1817–1827. doi:https://doi.org/10.1016/j.str.2008.10.008.

86. Sethi R, Seppälä J, Tossavainen H, Ylilauri M, Ruskamo S, Pentikäinen OT, et al. A Novel Structural Unit in the N-terminal Region of Filamins. Journal of Biological Chemistry. 2014;289(12):8588–8598. doi:https://doi.org/10.1074/jbc.M113.537456.

87. Papanikolopoulou K, Teixeira S, Belrhali H, Forsyth VT, Mitraki A, van Raaij MJ. Adenovirus Fibre Shaft Sequences Fold into the Native Triple Beta-Spiral Fold when N-terminally Fused to the Bacteriophage T4 Fibritin Foldon Trimerisation Motif. Journal of Molecular Biology. 2004;342(1):219–227. doi:https://doi.org/10.1016/j.jmb.2004.07.008.

88. Xiong X, Bromley EHC, Oelschlaeger P, Woolfson DN, Spencer J. Structural insights into quinolone antibiotic resistance mediated by pentapeptide repeat proteins: conserved surface loops direct the activity of a Qnr protein from a Gram-negative bacterium. Nucleic Acids Research. 2011;39(9):3917–3927. doi:10.1093/nar/gkq1296.

89. Coleman MA, Eisen JA, Mohrenweiser HW. Cloning and Characterization of HARP/SMARCAL1: A Prokaryotic HepA-Related SNF2 Helicase Protein from Human and Mouse. Genomics. 2000;65(3):274–282. doi:https://doi.org/10.1006/geno.2000.6174.

90. Schneidman-Duhovny D, Hammel M, Sali A. FoXS: a web server for rapid computation and fitting of SAXS profiles. Nucleic acids research. 2010;38:W540–W544.

91. Hura G, Menon A, Hammel M, Rambo R, Poole F, Tsutakawa S, et al. Robust, high-throughput solution structural analyses by small angle X-ray scattering (SAXS). Nature methods. 2009;6:606–12. doi:10.1038/nmeth.1353.

92. Svergun DI, Petoukhov MV, Koch MHJ. Determination of Domain Structure of Proteins from X-Ray Solution Scattering. Biophysical Journal. 2001;80(6):2946–2953. doi:https://doi.org/10.1016/S0006-3495(01)76260-1.

93. Boghog WU. Rossmann fold; 2022. Available from: https://en.wikipedia.org/wiki/Rossmann_fold#/media/File:Rossmann_fold_schematic.svg.

94. Rao ST, Rossmann MG. Comparison of super-secondary structures in proteins. Journal of Molecular Biology. 1973;76(2):241–256. doi:https://doi.org/10.1016/0022-2836(73)90388-4.

95. Hanukoglu I. Proteopedia: Rossmann fold: A beta-alpha-beta fold at dinucleotide binding sites. Biochemistry and Molecular Biology Education. 2015;43(3):206–209. doi:https://doi.org/10.1002/bmb.20849.

96. Vijayabaskar MS, Vishveshwara S. Insights into the Fold Organization of TIM Barrel from Interaction Energy Based Structure Networks. PLoS computational biology. 2012;8:e1002505. doi:10.1371/journal.pcbi.1002505.

